# MuseDrift: Navigating protein evolutionary manifolds with conditional discrete diffusion

**DOI:** 10.64898/2026.05.11.724439

**Authors:** Chaoyang Wang, Yiquan Wang

## Abstract

Protein engineering often requires variants that differ from a wild-type (WT) sequence by a specified amount while remaining structurally and functionally plausible. Controlling WT similarity alone does not specify which residues should change or how their substitutions should be coordinated. Here, we introduce MuseDrift, a conditional discrete diffusion model that generates variants from a WT sequence and a target sequence identity, without task-specific property guidance. Trained on WT–homolog pairs spanning different identities, MuseDrift uses pretrained WT representations as context and identity-aware modulation to condition denoising. Compared with editing baselines at the same requested identities, its variants achieve higher mean structural prediction confidence and greater similarity to predicted WT structures. On the CAMEO dataset, it tracks requested identities spanning 30%–95%, with mean absolute errors of 0.52–2.08 percentage points across targets. Its generated variants also receive more favorable scores from external mutation-effect predictors than those from the evaluated baselines. Together, these results support MuseDrift as a framework for controlled exploration of WT-centered protein sequence space.

## 1 Introduction

Protein engineering often begins with a known wild-type (WT) sequence whose structure or function provides a starting point for design. Natural variation and laboratory directed evolution motivate exploring sequence neighborhoods around this reference to improve stability, activity, or specificity (Arnold, 1998). The extent of change matters: conservative variants restrict exploration, whereas distant variants may lose the properties of the starting protein. Controllable WT-centered generation therefore aims to produce variants that balance proximity to the reference with diversity for further exploration (Figure 1A).

**Figure 1:**
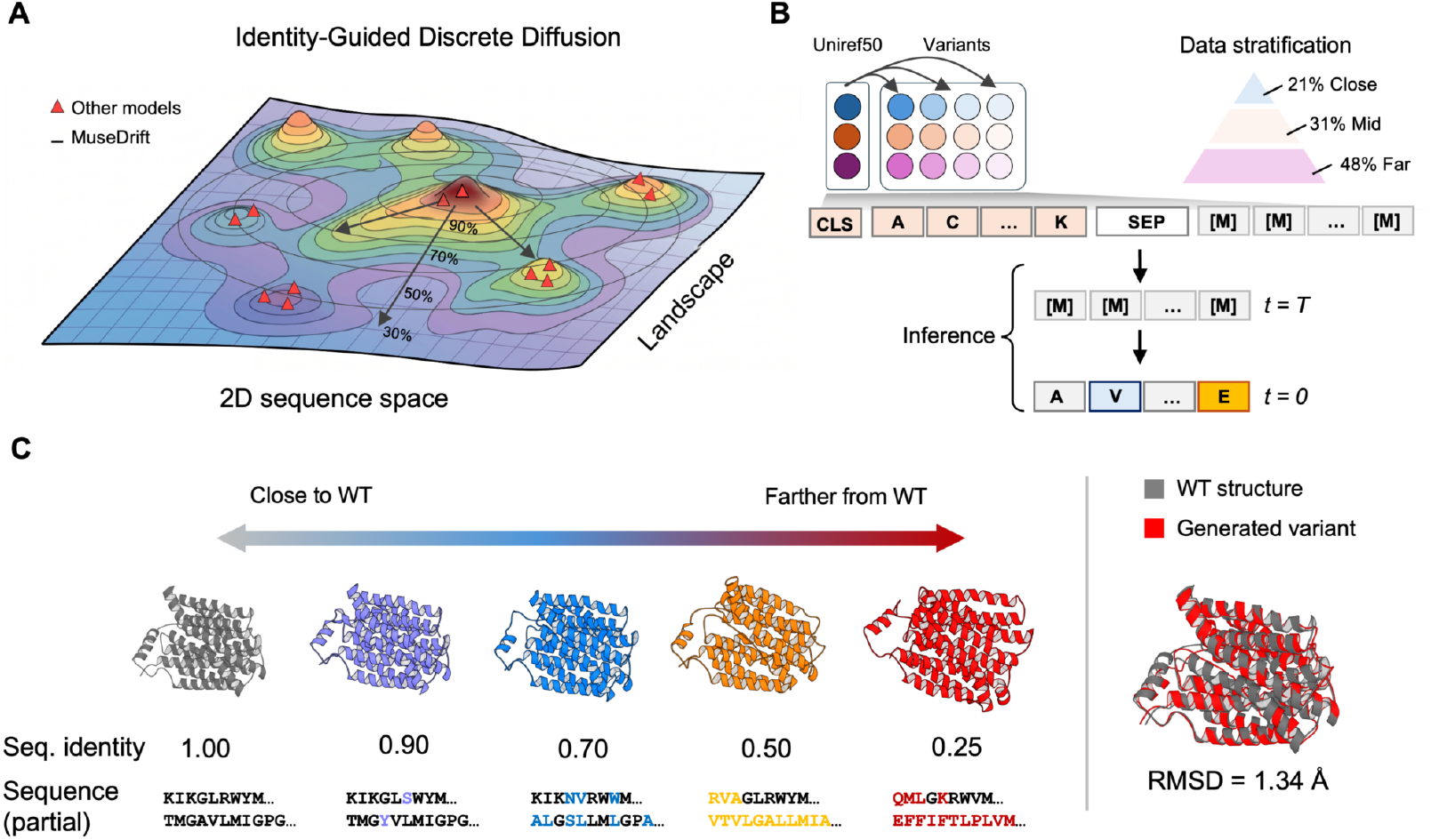
WT-centered protein variant generation. **(A)** Conceptual exploration around a reference protein, with lower target identity allowing greater sequence variation. **(B)** Seed-and-Stratify constructs 38.2M aligned WT–homolog pairs organized by identity. The schematic shows WT-prefix conditioning and iterative unmasking; *t* denotes denoising time. The model additionally uses frozen ESM-2 WT representations and identity FiLM. **(C)** Illustrative development-stage sequences and predicted structures at the displayed WT identities. The overlay reports the RMSD for the shown WT–variant pair. Quantitative results for the identity-conditioned model appear in Figure 2 and Table 1.

Recent protein language models and diffusion-based generators have advanced de novo and conditional protein design (Madani et al., 2023; Alamdari et al., 2023; Wang et al., 2024). Conditioning interfaces include sequence context, structural information, and descriptions of desired function (Hayes et al., 2025; Liu et al., 2025). WT-editing and evolutionary models also support reference-based exploration and mutation-burden control (Gruver et al., 2023; Deutschmann et al., 2026; Koehl et al., 2026). For WT-centered engineering, we investigate whether a generator learned from natural WT–homolog pairs can combine accurate identity control with favorable structural proxy scores. Generation is conditioned on the WT and requested identity, without externally prescribed mutation sites or task-specific property guidance.

We address this problem with MuseDrift, a conditional discrete diffusion model for WT-anchored protein generation. We use *evolutionary manifolds* to describe WT-centered neighborhoods shaped by natural substitution and conservation patterns. Seed-and-Stratify organizes WT–homolog pairs across identities, and the denoiser combines visible WT tokens, frozen ESM-2 WT representations, and direct numerical identity conditioning through FiLM (Figure 1B). Random-order iterative unmasking generates variants conditioned on the requested identity. Our evaluations show that MuseDrift combines structural quality, controllability, diversity, and favorable external scores, providing promising candidate pools for experimental screening.

### Contributions

1. We study WT-anchored generation as controlled exploration of sequence neighborhoods, using requested identity to specify distance while allowing the model to choose mutation sites and replacement residues.
2. We develop Seed-and-Stratify to construct 38.2M aligned WT–homolog pairs across 486K WT identifiers, organized by identity and split by WT identifier for conditional generation.
3. We introduce a conditional discrete diffusion model that combines WT-prefix context, frozen WT features, and identity FiLM with random-order unmasking for direct control of WT sequence identity.
4. We demonstrate accurate identity control and favorable structural proxy scores against the evaluated WT-conditioned controls, with smaller structural differences near the WT (Figure 2; Table 1).
5. We characterize library diversity, evolutionary substitution and mutation-placement patterns, and external fitness predictions (Figure 3A–D; Table 2; Appendix K), distinguishing computational evidence from experimentally established function.

## 2 Related Work

### Conditional protein generation

Protein generators condition on functional descriptions, sequence context, and structures. ProDVa augments a protein language model with a dynamically retrieved vocabulary of protein fragments for description-guided generation (Liu et al., 2025). ESM3 integrates sequence, structure, and function within a multimodal masked-generation framework (Hayes et al., 2025); its sequence-completion interface also supports sequence-only editing. WT-centered generation specifies a reference protein and target identity, making sequence distance an explicit input.

**Figure 2:**
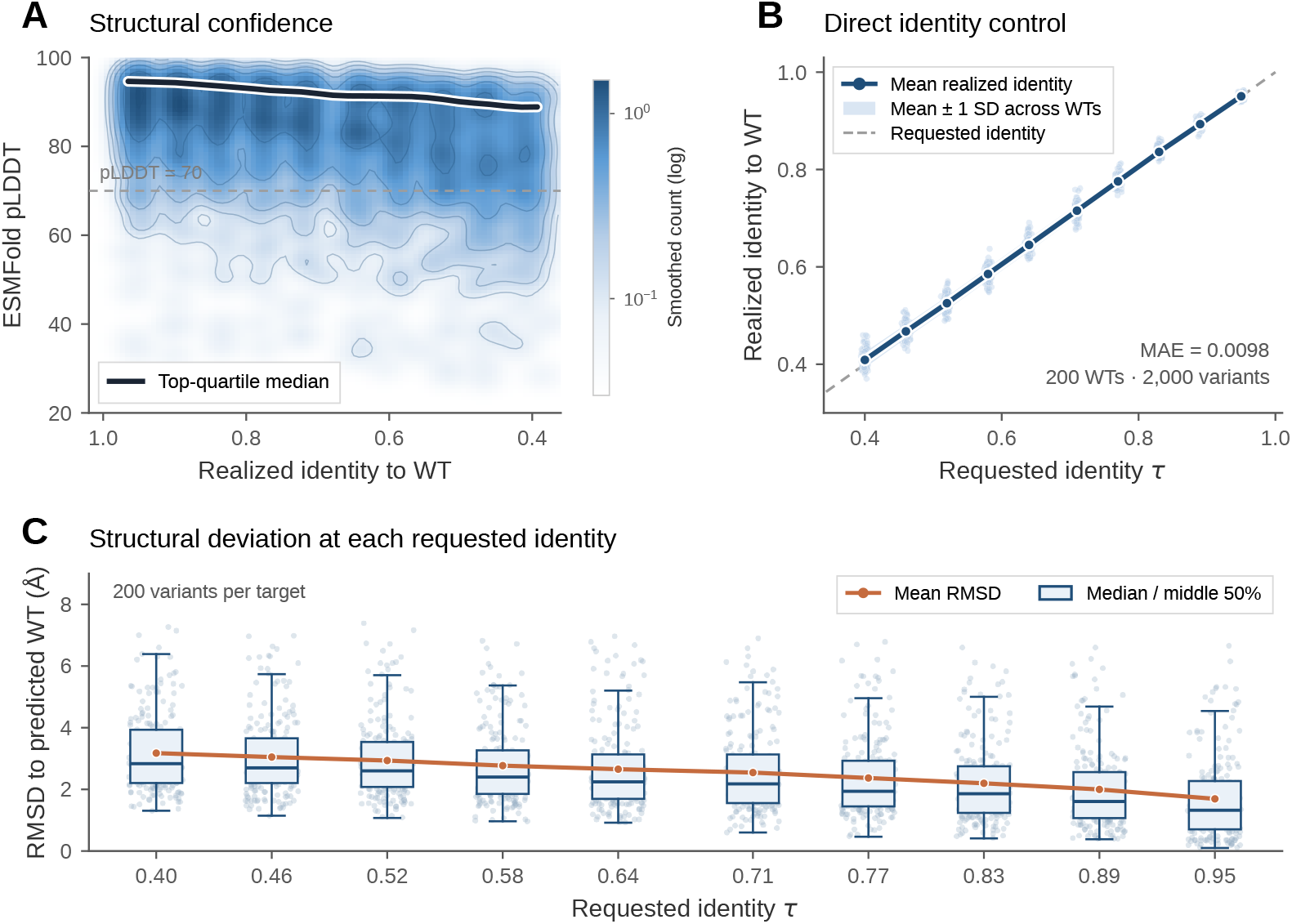
Identity-conditioned generation on the internal test set. All panels retain the same 2,000 variants: one per WT for 200 WTs at ten requested identities spanning .40–.95, with no quality-based selection. **A:** Smoothed density of realized identity and ESMFold pLDDT. The curve follows the smoothed median of the highest-pLDDT quartile within identity bins; the dashed line marks pLDDT 70. **B:** Requested versus realized identity: jittered individual points, mean curve, and one-standard-deviation ribbon across WTs; dashed line, *y* = *x*. **C:** RMSD of TM-align-aligned C*α* pairs between predicted variant and WT structures, grouped by requested identity. Points include all variants; boxes show medians and interquartile ranges with 1.5-IQR whiskers. The orange line joins means; lower RMSD indicates closer alignment.

**Figure 3:**
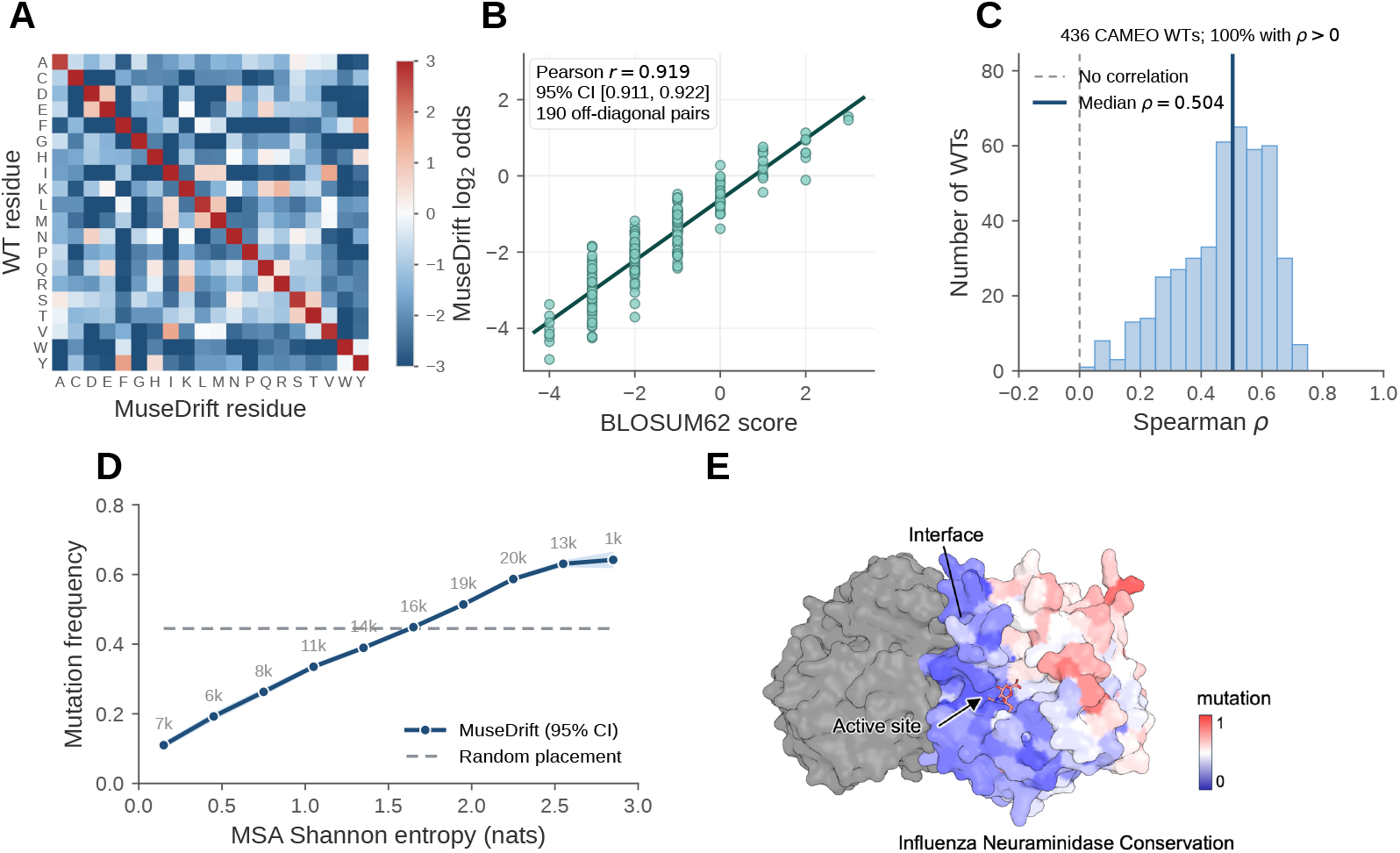
Evolutionary patterns in CAMEO generations (*τ* = .55, *K* = 8; A–D), with a structural reference illustration (E). **A:** WT-to-variant residue log_2_ odds on all 438 CAMEO WTs, including self-matches on the diagonal; colors saturate at ±3. **B:** BLOSUM62 scores versus symmetrized off-diagonal log odds (190 residue pairs); the interval uses 20,000 WT-hash Bayesian-bootstrap draws. **C:** Within-WT Spearman correlations between MSA entropy and mutation frequency on 436 CAMEO WTs with sufficient MSA coverage; two WTs lack valid entropy columns. Dashed and solid lines mark zero and the median. **D:** Mutation frequency averaged within each WT and entropy bin, then equally across WTs represented in that bin. Labels give the number of positions in thousands. The dashed line is the expectation under uniform random placement preserving each variant’s actual edit count. Shading shows 95% intervals from 20,000 WT-bootstrap resamples. **E:** Reference visualization of influenza neuraminidase with the active site and interface annotated. The mutation map ranges from blue (0) to red (1). This panel is a qualitative illustration.

### Reference-conditioned editing and evolution

EvoFlows learns edit flows between evolutionarily related proteins, jointly choosing edit locations and operations with a controllable number of edits (Deutschmann et al., 2026). PEINT learns sequence transitions conditioned on evolutionary time from phylogenetic sequence pairs, using frozen ESM representations (Koehl et al., 2026). DPLM-Evo predicts substitutions, insertions, and deletions during denoising to support variable-length evolution and post-editing (Wang et al., 2026). MuseDrift focuses on fixed-length variant generation with final sequence identity as the requested condition, rather than evolutionary time or an edit trajectory. Joint site and residue selection is shared with related approaches; our evaluation concerns the quality and identity response of this directly conditioned generator.

### Masked modeling and discrete diffusion

ESM-2 predicts masked residues from visible sequence context (Lin et al., 2023), allowing a pretrained model to fill selected positions in a WT. Discrete diffusion generalizes denoising to categorical variables (Austin et al., 2021), and EvoDiff applies iterative denoising to protein generation, including completion around fixed sequence segments (Alamdari et al., 2023). DPLM likewise learns protein generation through discrete diffusion (Wang et al., 2024). For WT editing, the fraction of masked positions does not by itself determine realized sequence identity: a predicted residue can reproduce the original WT residue. Enforcing substitutions at selected sites gives an explicit edit budget, but site selection then remains part of the sampling protocol. MuseDrift learns soft identity conditioning from WT–homolog pairs and generates variants by random-order unmasking (Bond-Taylor et al., 2022).

### Evaluation of WT-centered generation

The Continuous Automated Model EvaluatiOn (CAMEO) platform provides automated blind assessment of protein structure predictions (Haas et al., 2018). ProDVa adapts CAMEO targets into a function-conditioned generation benchmark by pairing reference proteins with function keywords (Liu et al., 2025). We use its sequence panel as WT references and assess identity control, structural quality, library diversity, evolutionary patterns, and mutation-effect predictions (§4.1). These computational readouts support candidate screening; retained function requires experimental validation.

## 3 Methods

MuseDrift generates variants by iterative masked-token denoising, conditioned on WT tokens, frozen ESM-2 features, and target identity.

### 3.1 Problem setup

Let *W* ∈ Σ*^L^* be a reference protein, denoted the WT, where Σ contains the 20 standard amino acids. Given *W* and a target identity *τ* ∈ (0, 1), the task is to sample variants *V_k_* ∼ *p_θ_*(*V* | *W, τ*) for *k* = 1*, …, K*. Each variant has the same length as the WT and permits substitutions only. Its realized identity is

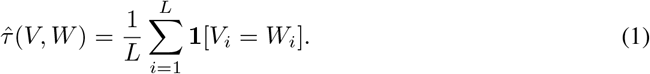

The aim is for *τ*^(*V, W*) to be close to *τ* (Figure 1A). Identity is a learned soft condition, so realized edit counts can vary between samples. We evaluate this control alongside the structural quality and diversity of the resulting libraries.

### 3.2 Seed-and-Stratify data pipeline

Seed-and-Stratify organizes natural WT–homolog pairs across identities to cover modest and extensive variation (Figure 1B). UniRef50 cluster representatives serve as seeds for sharded MMseqs2 searches against UniRef100 (Suzek et al., 2015; Steinegger & Söding, 2017). Retrieved homologs are divided into Far (40–70%), Mid (70–90%), and Close (90–100%) identity bins. Coverage filters remove short fragment matches, while per-WT, per-bin caps limit the influence of large homolog neighborhoods.

Each retained example is a triplet (*W*_aln_*, V*_0_*, τ*_train_): two locally aligned sequences, which may contain gaps, and their MMseqs2 local-alignment identity. Training receives numerical *τ*_train_, not bin labels; generation supplies requested full-length identity *τ* through the same pathway. Local-alignment identity differs from Eq. 1, so we evaluate this transfer by measuring requested versus realized identity (Appendix C).

The corpus contains approximately 38.2M pairs across 486K WT identifiers. Pairs are split 80/10/10 by WT identifier; homologous families may span splits. After applying the model’s context-length limit, approximately 23.3M training pairs over 337K WTs remain eligible. Exact filters, length limits, and split counts are given in Appendix A.

### 3.3 Conditional discrete denoising

#### Input and corruption

Let *V*_0_ be the uncorrupted aligned variant of length *L_V_*. We independently draw a corruption level *t* ∼ Uniform[0, 1) and replace max(1, ⌊*tL_V_* ⌋) randomly selected positions with `[MASK]`, producing *V_t_*. The model input is `[CLS]` *W*_aln_ `[SEP]` *V_t_*, with the WT prefix always visible. The corruption level *t* sets the mask count, while *τ*_train_ remains the observed identity of the uncorrupted pair. There is no separate *t* embedding; the mask pattern provides the corruption context.

#### Objective

The denoiser reconstructs masked variant tokens using the WT, visible variant context, and pair identity. For a batch B, with masked positions **m***_b_* for example *b*, the loss is

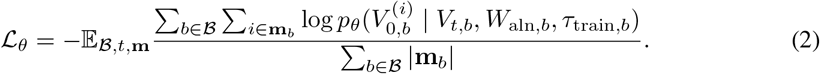

For brevity, Eq. 2 suppresses the full-WT feature input described in §3.4. This masked-token cross-entropy assigns equal weight to masked positions within a batch (Austin et al., 2021; Alamdari et al., 2023; Wang et al., 2024). The observed *τ*_train_ is held fixed during corruption and reconstruction; gradients from this loss train the backbone, WT adapter, and FiLM layers. Reconstructed residues may match or differ from the WT, so the corruption pattern specifies prediction targets rather than mutation sites. Further objective details are in Appendix B.

### 3.4 Conditioning architecture

A bidirectional Pre-LayerNorm transformer combines WT conditioning at the input with block-wise identity modulation.

#### WT context

The visible WT prefix is augmented with frozen full-WT ESM-2 t12 representations (Lin et al., 2023). A learned adapter applies LayerNorm and a linear projection to the denoiser width, then adds these features to WT-prefix embeddings before the transformer stack. Self-attention provides variant positions with this combined context. Feature assignment for local-alignment training and full-length generation is specified in Appendix C.

#### Identity FiLM

A sinusoidal embedding of the scalar identity condition and a two-layer MLP produce a scale and shift for each transformer block (Perez et al., 2018). For the feed-forward output *h_ℓ_* at block *ℓ*, conditioning takes the form

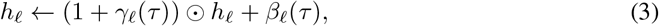

before dropout and residual addition, where *γ_ℓ_*(*τ*)*, β_ℓ_*(*τ*) ∈ R^768^. The modulation is shared across positions within each block.

The denoiser is trained from scratch, jointly optimizing the backbone, FiLM layers, and WT adapter, with 95.0M trainable parameters in total. The 33.5M-parameter ESM encoder remains frozen. Backbone dimensions, initialization, and optimization settings are given in Appendix B.

### 3.5 Identity-conditioned generation

Given a full-length WT and requested *τ*, we initialize a same-length, fully masked variant slot. WT context and *τ* remain fixed during decoding (Figure 1B; §3.4).

At each step, the denoiser predicts amino-acid distributions using the WT and the variant residues revealed so far. With *r* masks and *s* steps remaining, we reveal max(1, ⌊*r/s*⌋) uniformly selected positions and sample their residues from the softmax restricted to the 20 standard amino acids. Positions revealed in the same step are sampled independently from one forward pass, conditional on the shared WT and current variant context. Decoding uses at most 50 random-order reveal steps (Bond-Taylor et al., 2022) and stops when no masks remain. The reveal order specifies when positions are decoded; the sampled residues determine which positions ultimately differ from the WT.

Repeating this procedure with independent sampling seeds produces variants for the same (*W, τ*), with *τ* supplied directly to the learned conditioning pathway. No quality-based rejection is used; generation settings are given in Appendix B. For *L_V_ >* 1, training leaves at least one variant token visible, so the fully masked generation start extends beyond the training corruption support (Appendix C).

## 4 Experiments

### 4.1 Experimental setup

#### Evaluation panels

The main benchmark uses the CAMEO sequence panel released with ProDVa (Liu et al., 2025). Filtering to length *L* ≤ 511 retains 441 reference records corresponding to 438 unique WTs. Each main-grid method generates *K* = 64 variants per record at *τ* ∈ {.30, .40, .55, .75, .95}, giving 141,120 draws per method. Duplicate-record summaries are averaged within each WT before equal-WT aggregation. Identity and diversity use all 64 samples per record; structural comparisons use the predesignated first sample. A separate *K* = 8 draw on the same references supports substitution and conservation analyses; 436 WTs meet the MSA coverage criteria. External fitness scoring uses all samples on 429 shared WTs (Appendix O). CAMEO is not family-held-out; training overlap and panel definitions are detailed in Appendices J and D.

#### Baselines

WT-conditioned baselines receive sequence and target identity; structures are used only for evaluation. Four main-grid exact-edit controls introduce *n_τ_* (*L*) = round((1−*τ*)*L*) substitutions. For each (*W, τ, k*), they share a seeded edit-site set, resampled for each sample index *k*. Uniform draws replacement residues uniformly, while BLOSUM62-odds weights them by substitution odds. ESM-2 full-mask uses a visible WT prefix and a fully masked variant slot, decoded in up to 50 random-order reveal steps. EvoDiff OA-DM fills selected sites within the WT. All four forbid the WT residue at edit sites. ESM-2 selected-site infilling leaves unedited WT residues visible in place and fills masked sites in one forward pass with non-WT residues. Both ESM-2 protocols use the same frozen 650M model. Infilling covers all 438 WTs and five targets with *K* = 8; structure scoring uses the first sample. MuseDrift chooses its own sites and replacement residues. These are comparisons of complete generation procedures; they do not separate the contributions of site selection and residue sampling. Full protocols are in Appendices G and H.

#### Metrics

Structural proxies are single-sequence ESMFold pLDDT (Lin et al., 2023) and TM-align similarity between predicted variant and predicted WT structures, normalized by WT length (Zhang & Skolnick, 2005). Neither score is used for sample selection. Identity error measures deviation from the request; library diversity uses sequence uniqueness, pairwise identity, and edit-set Jaccard distance. Unless stated otherwise, results are equal-WT means, with 95% intervals obtained by resampling WTs and preserving pairing between methods (Appendix D).

### 4.2 Structural quality

#### Structural response to the identity dial

Figure 2A relates predicted structural confidence to realized identity on the internal test panel. Mean aligned RMSD to the predicted WT decreases from 3.18 Å at *τ* = .40 to 1.69 Å at .95 (Figure 2C). Both panels retain all 2,000 variants; RMSD uses TM-align-aligned residues.

#### CAMEO benchmark

MuseDrift has the highest mean pLDDT and predicted-WT-relative TM-score at all five targets among the methods in Table 1. Its mean pLDDT increases from 66.46 at *τ* = .30 to 84.28 at .95, while predicted-WT-relative TM-score increases from .712 to .940. At *τ* = .30*/.*40, EvoDiff is the strongest external baseline; MuseDrift exceeds its pLDDT by 26.40/26.82 and its TM-score by .334/.309. ESM-2 infilling reaches pLDDT 33.90/35.18 at these targets.

ESM-2 selected-site infilling is the strongest external baseline at *τ* = .55*/.*75*/.*95. At *τ* = .55*/.*75, MuseDrift exceeds its pLDDT by 8.33 [7.19, 9.49] and 2.48 [1.93, 3.04], respectively. Near the WT at *τ* = .95, the pLDDT difference is +.21 [−.04, .46], while the TM-score difference remains positive at +.010 [.004, .017]. The advantage over these baselines is clearest at larger sequence changes and narrows near the WT. Full-mask and infilling differ in both input layout and decoding, so their difference does not isolate the effect of visible WT context (Appendix H).

#### Filtered CAMEO results

After excluding exact training matches and detected matches with ≥ 50% identity and ≥ 80% coverage of both sequences, 223 CAMEO WTs remain. On this subset, MuseDrift retains the highest mean pLDDT and predicted-WT-relative TM-score at all five target identities (Appendix J, Table 9).

**Table 1:** Structural quality of WT-conditioned CAMEO generations. Equal-WT means on 438 WTs, using the predesignated first variant per method and identity setting. Bold marks the highest point estimate; paired intervals and realized identities are in Appendices F, H, and Q.

| Method | Parameters | Target identity $\tau$ | | | | |
| --- | --- | --- | --- | --- | --- | --- |
|  |  | .30 | .40 | .55 | .75 | .95 |
| *ESMFold pLDDT* ↑ | | | | | | |
| MuseDrift | 95M<sup>†</sup> | **66.46** | **71.32** | **76.29** | **81.24** | **84.28** |
| ESM-2 infilling | 650M | 33.90 | 35.18 | 67.96 | 78.76 | 84.06 |
| ESM-2 full-mask | 650M | 32.71 | 33.19 | 47.28 | 72.70 | 83.20 |
| EvoDiff OA-DM | 38M | 40.06 | 44.50 | 54.34 | 70.35 | 83.03 |
| BLOSUM62-odds | — | 30.93 | 35.33 | 51.92 | 71.32 | 82.94 |
| Uniform | — | 29.51 | 30.07 | 39.73 | 66.61 | 82.42 |
| *Predicted-WT-relative TM-score* ↑ | | | | | | |
| MuseDrift | 95M<sup>†</sup> | **0.712** | **0.764** | **0.812** | **0.866** | **0.940** |
| ESM-2 infilling | 650M | 0.261 | 0.330 | 0.727 | 0.841 | 0.930 |
| ESM-2 full-mask | 650M | 0.272 | 0.304 | 0.539 | 0.805 | 0.920 |
| EvoDiff OA-DM | 38M | 0.378 | 0.455 | 0.594 | 0.773 | 0.913 |
| BLOSUM62-odds | — | 0.294 | 0.379 | 0.610 | 0.792 | 0.922 |
| Uniform | — | 0.269 | 0.290 | 0.455 | 0.764 | 0.917 |
| *Internal configuration: WT-token-only* |  |  |  |  |  |  |
| pLDDT | 85M | 46.40 | 56.13 | 64.74 | 75.49 | 83.12 |
| TM-score | 85M | 0.520 | 0.620 | 0.715 | 0.826 | 0.936 |

<sup>†</sup>Trainable parameters, plus a frozen 33.5M ESM-2 WT encoder. WT-token-only is a separately trained, temperature-calibrated configuration; training schedules and realized identities differ from the full model (Appendix Q). Bold does not indicate statistical significance.

#### Conditioning configurations

We compare the full model with three trained configurations that omit identity FiLM, ESM-2 WT features, or both. All configurations retain WT tokens. At nominal identity .55, mean pLDDT is 76.29 for the full model, 72.21 without ESM-2 features, 71.85 without FiLM, and 64.74 for the WT-token-only configuration. The corresponding realized identities are .555/.553/.530/.502. All four use the same 438 WTs. Differences in training schedules, calibration, and realized identity prevent causal attribution to individual components (Appendix Q, Table 21).

#### Energy-based assessment

On a separate 80-WT CAMEO subset, FoldX (Schymkowitz et al., 2005) reports lower length-normalized energy changes for MuseDrift than Uniform, BLOSUM62-odds, and EvoDiff at all four targets (Figure 4A,B). Against ESM-2 infilling, the difference at *τ* = .55 is −.209 [−.282, −.141] kcal/mol/residue; intervals include zero at *τ* ≥ .75. These energies are computed on ESMFold structures and complement the confidence scores (Appendix M).

#### Scope of the comparison

Realized identities are close but not identical between MuseDrift and the external controls. Appendix E examines sensitivity to these differences for the four main-grid controls and ESM-2 selected-site infilling. All structural and energy scores are computational assessments.

### 4.3 Controllability and diversity

#### Identity control

Figure 2B shows that realized identity follows the request on the internal test panel, with overall mean absolute error .0098. On CAMEO, the *K* = 64 mean absolute identity errors are .0208/.0155/.0131/.0111/.0052 across the five targets (Table 7). The .30 request is below the .40 lower bound of training-label values; the other four requests fall within that numerical range. Because training uses local alignments, all five also test transfer to full-length generation. The fraction within .05 of the request rises from 94.1% at *τ* = .30 to 99.4% at .95. This response is obtained by changing the identity input alone. Exact-edit controls achieve their requested counts by construction, leaving only integer-rounding error. Short references and targets near one expose limits of aggregate control; per-WT results and a high-identity extension are in Appendix I.

#### Library diversity

All 2,205 MuseDrift reference-record–target libraries contain 64 distinct sequences and no exact WT copies (Appendix K, Table 11). Mean pairwise identity rises from .231 at *τ* = .30 to .908 at .95, reflecting narrower libraries nearer the WT. Compared with the random-site controls, MuseDrift has higher pairwise identity and lower edit-set Jaccard distance. At *τ* = .55, mean pairwise identity is .418 versus .313–.323 for the controls; paired WT-bootstrap intervals support these differences (Appendix K, Table 12). Each target therefore yields distinct variants, but with less diversity than these controls. The comparison does not establish a causal quality–diversity trade-off.

### 4.4 Evolutionary patterns and predicted fitness

#### 4.4.1 Evolutionary characterization

##### Substitution preferences

On all 438 CAMEO WTs, MuseDrift’s pooled substitution log-odds correlate with BLOSUM62 (Henikoff & Henikoff, 1992) at *r* = .919*/.*863*/.*693*/.*592 for *τ* = .55*/.*75*/.*90*/.*95 (Figure 3A,B), consistent with evolutionary substitution preferences. ESM-2 infilling and EvoDiff also show positive correlations, while BLOSUM62-odds uses the reference matrix directly (Appendix N).

##### Mutation placement

Across 436 CAMEO WTs with sufficient MSA coverage, median within-WT Spearman correlations between MSA entropy and mutation frequency are .504/.436/.313/.234 at *τ* = .55*/.*75*/.*90*/.*95, and 97.0–100% of WTs have positive correlations (Figure 3C). At *τ* = .55, mutation frequency rises with entropy, unlike the approximately flat uniform-placement reference matched by actual edit counts (Figure 3D; Appendix N). Thus, variants preferentially mutate less-conserved positions.

**Table 2:** External predicted fitness of generated libraries on 429 CAMEO WTs. WT-relative additive scores (WT = 0; higher is better), averaged over all generated samples within each WT and then equally across WTs. Main-grid methods use *K* = 64; ESM-2 infilling uses *K* = 8. Bold marks the highest mean per column within each predictor. Scores are predictions, not measured fitness. Pair<u>ed</u> intervals and sensitivity analyses are in Appendix O.

| Generator | Target identity $\tau$ | | | | |
| --- | --- | --- | --- | --- | --- |
|  | .30 | .40 | .55 | .75 | .95 |
| *ProSST-2048* ↑ | | | | | |
| MuseDrift | **-732.11** | **-603.91** | **-429.10** | **-222.38** | **-43.38** |
| ESM-2 infilling | -1149.26 | -939.74 | -538.20 | -275.61 | -53.91 |
| ESM-2 full-mask | -1129.00 | -959.08 | -687.89 | -344.00 | -64.33 |
| EvoDiff OA-DM | -1045.49 | -867.92 | -625.32 | -334.14 | -64.96 |
| BLOSUM62-odds | -1024.36 | -878.46 | -658.49 | -365.72 | -73.38 |
| Uniform | -1141.13 | -978.10 | -733.15 | -407.46 | -81.75 |
| *VenusREM* ↑ | | | | | |
| MuseDrift | **-167.91** | **-136.95** | **-96.23** | **-49.35** | **-9.55** |
| ESM-2 infilling | -268.33 | -220.33 | -127.63 | -65.57 | -12.84 |
| ESM-2 full-mask | -263.78 | -224.20 | -161.27 | -81.20 | -15.25 |
| EvoDiff OA-DM | -245.37 | -204.03 | -147.31 | -78.90 | -15.37 |
| BLOSUM62-odds | -241.83 | -207.36 | -155.44 | -86.33 | -17.32 |
| Uniform | -266.63 | -228.52 | -171.30 | -95.20 | -19.10 |

#### 4.4.2 External mutation-effect assessment

We score the generated libraries with ProSST-2048 (Li et al., 2024) and VenusREM (Tan et al., 2025), which combines ProSST-2048 with MSA information. Both use the same predicted WT structure across generators. Scores sum WT-relative substitution log odds; higher values indicate more favorable predictions, with the WT assigned zero. We average all samples (*K* = 64 for main-grid methods; *K* = 8 for ESM-2 infilling) within WT, then equally across 429 WTs, without score-based selection.

Among the evaluated generators, MuseDrift has the highest mean score under both predictors at every target (Table 2). At *τ* = .55, its advantage over ESM-2 infilling is 109.11 [102.49, 115.84] under ProSST-2048 and 31.41 [29.57, 33.29] under VenusREM. The advantage persists when all methods use the first eight samples, when scores are normalized by each variant’s actual substitution count, and with ProSST-4096 (Appendix O). VenusREM shares the ProSST-2048 component, so the assessments are complementary rather than independent. The additive scores do not model interactions among mutations, and higher predicted scores do not establish experimentally retained function.

## 5 Conclusion and Limitations

MuseDrift generates distinct WT-centered variants with direct control of sequence identity. Structural proxy scores exceed those of the evaluated exact-edit controls, with smaller differences from ESM-2 infilling near the WT. Generated substitutions follow evolutionary preferences, and external mutation-effect predictors favor its libraries.

Our evaluation relies on computational predictions, so experimental validation remains necessary to establish the activity and engineering utility of generated variants. The current model conditions on sequence context and target identity; future work will incorporate explicit structural information to guide variant generation under structural constraints.

### AI USE STATEMENT

Generative AI tools assisted with literature organization, evaluation-protocol review, experimental and analysis-code development, inspection and interpretation of computational results, figure preparation, and drafting and revising manuscript text. Reference sequences and experimental assay labels came from the stated data sources. The authors are responsible for verifying the experimental protocols, numerical results, citations, and final scientific claims.

## Ethics statement

This computational study uses public protein sequence and assay data and involves no human subjects or personal data. Protein variant generation has beneficial research applications and potential for misuse. The reported computational measurements do not establish activity, viability, pathogenicity, or safety; experimental use requires appropriate biosafety review and empirical validation.

## Reproducibility statement

Sections 3 and 4 describe the model and evaluation. Appendices A–C describe data construction, optimization, generation, and implementation limitations. The remaining appendices give baseline protocols, sample scopes, uncertainty estimates, and auxiliary analyses. Configurations, available checkpoint and artifact checksums, sampling seeds, and evaluation scripts are recorded with the computational results. The historical WT-embedding cache lacks an authenticated upstream encoder revision.

## A Data construction and split details

### Source snapshots and construction settings

The UniRef100 database records release `2026 01` (28 January 2026). The UniRef50 seeds are a frozen REST export with query `identity:0.5 AND count:[50 TO *]`; its compressed FASTA has SHA-256 prefix `dd5162444abd`. The export does not retain a UniRef50 release tag. The 20-shard MMseqs2 search uses sensitivity 6, E-value threshold 10^−3^, minimum identity .40, target coverage .80 (`cov-mode 1`), and stored alignments (`-a`).

Seed-and-Stratify starts from UniRef50 representatives with at least 50 cluster members and searches UniRef100 with MMseqs2. Hits must have identity in [0.40, 1.00), at least 70% WT coverage and 80% variant coverage; self hits, identical aligned strings and duplicate identifier pairs are removed. For each WT, retained neighbors are split into 40–70%, 70–90% and 90–100% identity bins, capped at 67 per bin and 200 total. We then require at least three neighbors in every bin. These bin boundaries and caps were not tested in a matched sensitivity study, so we make no robustness claim for those balancing choices. The construction yields 38,198,695 aligned pairs over 486,037 WTs. Before applying the caps, each alignment shard is shuffled with pandas `random state=13`. The retained construction summary has SHA-256 prefix `bae1b3a0a3ba`.

WT identifiers, rather than individual pairs, are randomly assigned 80/10/10 to train/validation/test with seed 42, so all pairs for one WT remain in one split. Specifically, sorted WT identifiers are shuffled with NumPy `RandomState(42)` with *N* = 486,037; the first ⌊0.8*N* ⌋ and next ⌊0.1*N* ⌋ identifiers form train and validation, with the remainder assigned to test. The split manifest (SHA-256 prefix `3438124a0651`) records 30,557,294/3,826,390/3,815,011 pairs across 388,829/48,603/48,605 WT identifiers before length filtering. The model input [`CLS`, WT, `SEP`, Var] has length 2*L* + 2; with a 1,024-token limit, the loader retains only aligned length *L* ≤ 511. After applying this model-time filter, 29,167,112 pairs over 420,994 WTs are eligible: 23,342,731/336,856 for training, 2,911,314/42,097 for validation and 2,913,067/42,041 for test. These are loader-eligible split counts rather than a unique-row consumption ledger for the optimizer.

## B Architecture, optimization, and generation

MuseDrift uses 12 Pre-LayerNorm transformer blocks with rotary position embeddings (Su et al., 2021), width 768, and 12 attention heads; backbone and token-embedding dropout is 0.1, while the WT-adapter dropout is 0.0. The 480-dimensional frozen ESM-2 t12 WT representation is layer-normalized and projected once to width 768. A 256-dimensional sinusoidal identity embedding is mapped by a 256 → 512 → (2 × 12 × 768) MLP to per-block FiLM scale and shift parameters. The model has 95,018,203 trainable parameters; the frozen ESM-2 dependency has 33,500,401 parameters and is not stored in the MuseDrift checkpoint. The final FiLM projection and the WT-feature projection are zero-initialized. There is no separate embedding of the corruption level *t*; the input mask pattern provides the corruption context.

The training objective in Eq. 2 is masked-token cross-entropy, averaged over all masked variant tokens in a batch. WT positions and visible variant positions do not contribute to the loss. This objective does not implement the exact D3PM ELBO; no additional identity-error, structure, or fitness loss is used.

Training is from random initialization on eight NVIDIA B200 GPUs with bfloat16 automatic mixed precision. The per-GPU batch is 56 with two-step gradient accumulation, for an effective batch of 896 sequences. We use AdamW with *β* = (0.9, 0.98), peak learning rate 3 × 10^−4^, weight decay 0.01, 10,000 linear-warmup steps and cosine decay to 3 × 10^−5^ over at most 228,000 optimizer steps; gradient norm is clipped at 0.5. Validation is capped at 2,000 batches per epoch. Within the 228,000-step budget, we select the epoch-end checkpoint with the lowest recorded validation loss: epoch 8, step 208,416, with loss 1.037951 and SHA-256 prefix `031267dce3a7`.

The historical run records its hardware and optimization configuration but does not retain a complete, authenticated allocation-time ledger. We therefore make no runtime, cost, or parameter-efficiency comparison from this checkpoint. Unless stated otherwise, generation starts with an entirely masked variant slot and runs up to 50 random-order reveal steps, stopping when no masks remain. Sampling uses the standard softmax (temperature 1.0), without rescaling logits. Benchmark outputs are equal-length substitutions restricted to the 20 standard amino acids. The target identity is supplied directly to MuseDrift, without post-hoc identity calibration.

## C Training and generation details

### Sequence representation and identity labels

The stored pairs consist of MMseqs2 local aligned strings `qaln`/`taln`; their label is local-alignment percentage identity. The training vocabulary includes gap tokens. By contrast, the reported generation task starts from an unaligned full WT and emits a same-length string over the standard 20 amino acids. Consequently, the training label and output Hamming identity are different quantities, and the measured identity-response curve is an empirical transfer between these domains.

### WT feature assignment

Let *E*_0_*, …, E_L_*_−1_ denote the cached full-WT residue features. For a local WT segment with *m* non-gap residues, training assigns *E*_0_*, …, E_m_*_−1_ sequentially to its non-gap positions; gaps receive no ESM-2 features and do not advance the index. The stored aligned pairs do not retain the segment’s zero-based start coordinate *s*. For *s >* 0, the assigned feature indices therefore differ from the corresponding full-WT positions *s, …, s* + *m* − 1. Full-length generation assigns *E_i_* to WT residue *i*. The reported MuseDrift checkpoint was trained with this assignment; the effect of using residue-matched features during training has not been isolated.

### Corruption support and encoder provenance

For variant length greater than one, the rule max(1, ⌊*tL_V_* ⌋) with *t <* 1 never masks the complete variant. Inference nonetheless starts from a fully masked slot. The experiments measure the resulting implementation, but do not isolate this endpoint mismatch or establish that changing it would improve performance. The historical cache records the ESM-2 model identifier but lacks an authenticated upstream revision; this limits exact reconstruction of its original preprocessing environment. The reported model comparisons evaluate the combined conditioning architecture; gains from these comparisons alone cannot be attributed to individual conditioning paths.

### Internal-test illustration in **Figure 2**

We uniformly sample 200 full WTs without replacement, with seed 20260923, from 35,813 eligible unique sequences associated with test-split identifiers. The full-WT lookup is restricted to standard amino acids and length 60–511, then deduplicated. Exact matches to training WTs and the stored calibration references are excluded; selected lengths are 65–511. This identifier-based split does not exclude homologous families or overlap with training variant sequences. One variant is generated per WT at each of *τ* ∈ {.40, .46, .52, .58, .64, .71, .77, .83, .89, .95}, with all 2,000 outputs retained. Figure 2 uses single-sequence ESMFold predictions and TM-align RMSD over aligned C*α* pairs. The pLDDT density uses a smoothed 90 × 90 histogram; the upper-quartile curve uses 30 identity bins, at least 12 observations per bin, and two five-point moving averages. Identity dispersion is the sample standard deviation across WTs, not uncertainty across training runs. Panel C groups the same variants by requested identity. Boxes show the median and interquartile range, with 1.5-IQR whiskers; all individual RMSDs are plotted with horizontal jitter, and the line joins arithmetic means.

## D Evaluation panels and statistical protocols

**Table 3:** Evaluation panels and sample scopes. *K* is the number of generated samples per reference and target unless a scored subset is specified. Structural scores use ESMFold. Target grids and selection rules are defined below.

| Analysis | Reference source | Targets; samples | Analyzed members and selection |
| --- | --- | --- | --- |
| CAMEO structure, identity, diversity | ProDVa CAMEO panel; 441 records, 438 unique WTs | $T_C$; $K = 64$ per record | First sample for structure, all 64 for identity/diversity; length $\leq 511$; duplicate-record summaries averaged within WT. |
| Filtered CAMEO structure | Subset of the same panel; 226 records, 223 unique WTs | $T_C$; same first samples | Exclude exact matches and detected training hits with identity $\geq 50\%$, coverage $\geq 80\%$ on both sequences; audit both training-pair sides (Appendix J). |
| ESM-2 infilling structure | Same 438 CAMEO WTs | $T_C$; $K = 8$ | First sample only; selected-site, single-pass infilling. MuseDrift uses the main-grid first sample. |
| ProDVa structure | Fixed 429-WT CAMEO subset; 432 reference descriptions | No target; three draws per record | All 1,296 draws accounted for; Coverage and aggregate scores in Appendix R; no identity-band performance comparison. |
| FoldX energy | Frozen 80-WT CAMEO subset | $T_E$; one scored member | Predesignated sample 0 and its WT reference; separate ESMFold structures. ESM-2 uses selected-site infilling. |
| Substitution statistics | All 438 CAMEO WTs | $T_E$; $K = 8$ | All eight members, equal-WT aggregation; ESM-2 uses selected-site infilling. Separate generation draw from the main K64 grid. |
| MSA entropy | Same CAMEO panel; 436 of 438 unique WTs | $T_E$; $K = 8$ | Same draw as substitution statistics; at least 30 columns with $\geq 10$ homolog observations and nonconstant entropy. Two WTs lack valid columns. All eight variants per target. |
| External fitness prediction | Fixed 429-WT subset of the CAMEO panel | $T_C$; $K = 64 / K = 8$ | All main-grid / ESM-2 infilling samples; nine annotation-flagged references excluded before scoring. Same WT structures and MSAs across methods. |
| ProteinGym scoring | 162 single-mutant / 65 multi-mutant assays; 141 / 62 exact-WT clusters | $\tau = 1 - \vert M\vert /L$; no generation | At most 5,000 assayed variants per assay; standard-AA reference, $L \leq 511$, valid substitutions and finite DMS scores. |

The target grids in Table 3 are *T_C_* = {.30, .40, .55, .75, .95} and *T_E_* = {.55, .75, .90, .95}. The request .30 extrapolates below the numerical training-label range [.40, 1). It also changes the sequence representation and identity definition from local alignments to full-length substitutions, so it does not isolate numerical extrapolation from this domain transfer.

### CAMEO references

Of the 507 ProDVa CAMEO records, 441 have length at most 511; collapsing three duplicate-sequence pairs gives 438 unique references. The substitution analysis uses this entire panel, with a separate K8 draw. Conservation uses that same draw and query-matched MSAs on the CAMEO references. A valid entropy column requires at least ten standard-amino-acid homolog observations, excluding the query. Each WT must have at least 30 valid columns and non-constant entropy. Of 438 WTs, 436 meet these criteria; the remaining two have no valid columns. Their structural and substitution results are retained in the full-panel analyses. The 80-WT subset for FoldX was fixed before the comparison. It excludes a historical calibration list and references flagged by the training-WT-side exact/high-identity screen, then samples 80 evenly spaced ranks after sorting eligible records by length. It spans 38–507 residues. The subsequent training-variant-side screen flags seven of these references, so this subset is not a family-held-out or fully training-disjoint benchmark.

### Aggregation and uncertainty

Sample scores and library metrics are computed within each reference record, then duplicate-record summaries are averaged within each WT sequence; unique WT sequences receive equal weight. Library metrics are computed before this averaging, without pooling the samples of duplicate records. Method differences preserve pairing within WT. The CAMEO structural comparisons and library-diversity contrasts use 20,000 WT-bootstrap replicates; FoldX uses 10,000. Resampling holds the checkpoint and generated sequences fixed. These intervals do not quantify variation across training runs or separately estimate within-WT generation variability. They are unadjusted for multiple comparisons and do not account for homology between different WT sequences.

Substitution-matrix intervals use a WT-level Bayesian bootstrap, described in Appendix N. The MSA analysis reports within-reference correlations; its interquartile ranges describe the spread across references and are not confidence intervals. ProteinGym averages assay-level correlations and resamples exact-WT clusters, keeping assays of the same reference together.

## E Local-slope sensitivity to the model–control identity difference

MuseDrift’s identity conditioning is learned and therefore soft, so its realized identity differs slightly from the controls’ exact edit budget. The difference is positive through *τ* = 0.75 and slightly negative at 0.95. Table 4 estimates the corresponding signed shift for the four main-grid controls and ESM-2 selected-site infilling. For each control and target, we estimate its local pLDDT–identity slope from its neighboring targets and multiply by the realized identity difference. Interior targets use the adjacent lower and higher targets; endpoints use a one-sided difference. Slopes use realized identities and unrounded equal-WT means. For the original four controls, shift magnitudes are 0.2%–5.3% of the observed gaps. For ESM-2 infilling, the signed shifts at *τ* = .55*/.*75 are +.65*/* + .20 pLDDT, or 7.9%*/*8.2% of the corresponding gaps. These are local-linear diagnostics, not causal attributions or formal bounds. In particular, the small infilling gap at .95 has an interval spanning zero (Table 6).

## F Paired contrasts against random-site controls

Table 5 reports the paired structural-score differences.

## G Generation-control protocols

The four main-grid exact-edit controls share a position seed keyed by (*W, τ, k*), independent of method. Changing *k* resamples edit sites; replacement random streams are method-specific. They take *n* = round((1−*τ*)*L*) positions without replacement and forbid the WT residue at each selected site. Uniform samples from the other 19 residues. BLOSUM62-odds samples proportionally to 2*^Bij/^*^2^. For ESM-2 t33 650M, the WT is a visible prefix and the full variant slot begins masked; during at most 50 random-order reveals, unedited positions are forced to WT and edited positions to one of the other 19 amino acids. EvoDiff OA-DM instead masks only the selected sites in an otherwise visible WT and iteratively reveals non-WT residues. The controls receive this external random-site stream, while MuseDrift selects its own sites and residues; the comparison therefore does not separate those two sources of advantage.

**Table 4:** Local-linear sensitivity to realized identity. For each control, its pLDDT–identity slope between neighboring targets is multiplied by the model–control identity difference (Δid). Gap denotes MuseDrift minus control pLDDT. Calculations use unrounded equal-WT means for the pre-designated first sample on all 438 WTs; slopes are pLDDT per unit identity. Shift magnitudes are 0.2%–5.3% of gaps for the original four controls; ESM-2 infilling gives 7.9%*/*8.2% at .55*/.*75. This diagnostic is neither a causal attribution nor a formal bound.

| Control | $\tau$ | Gap | $\Delta\text{id}$ | Slope | Signed shift | $\vert \text{shift}\vert /\vert \text{gap}\vert $ |
| --- | --- | --- | --- | --- | --- | --- |
| ESM-2 650M full-mask | 0.3 | +33.76 | +0.0164 | 4.9 | +0.08 | 0.2% |
|  | 0.4 | +38.13 | +0.0075 | 58.3 | +0.44 | 1.1% |
|  | 0.55 | +29.01 | +0.0053 | 112.9 | +0.59 | 2.0% |
|  | 0.75 | +8.54 | +0.0050 | 89.9 | +0.45 | 5.3% |
|  | 0.95 | +1.07 | −0.0004 | 52.5 | −0.02 | 2.2% |
| EvoDiff OA-DM 38M | 0.3 | +26.40 | +0.0164 | 44.4 | +0.73 | 2.8% |
|  | 0.4 | +26.82 | +0.0075 | 57.1 | +0.43 | 1.6% |
|  | 0.55 | +21.95 | +0.0053 | 73.9 | +0.39 | 1.8% |
|  | 0.75 | +10.89 | +0.0050 | 71.8 | +0.36 | 3.3% |
|  | 0.95 | +1.25 | −0.0004 | 63.4 | −0.03 | 2.3% |
| BLOSUM62-odds | 0.3 | +35.53 | +0.0164 | 44.0 | +0.72 | 2.0% |
|  | 0.4 | +35.99 | +0.0075 | 83.9 | +0.63 | 1.8% |
|  | 0.55 | +24.37 | +0.0053 | 102.8 | +0.54 | 2.2% |
|  | 0.75 | +9.92 | +0.0050 | 77.6 | +0.39 | 3.9% |
|  | 0.95 | +1.34 | −0.0004 | 58.1 | −0.03 | 1.9% |
| Uniform | 0.3 | +36.95 | +0.0164 | 5.5 | +0.09 | 0.2% |
|  | 0.4 | +41.25 | +0.0075 | 40.8 | +0.31 | 0.7% |
|  | 0.55 | +36.56 | +0.0053 | 104.4 | +0.55 | 1.5% |
|  | 0.75 | +14.63 | +0.0050 | 106.8 | +0.54 | 3.7% |
|  | 0.95 | +1.86 | −0.0004 | 79.1 | −0.04 | 1.9% |
| ESM-2 infilling | 0.3 | +32.56 | +0.0164 | 12.8 | +0.21 | 0.6% |
|  | 0.4 | +36.14 | +0.0075 | 136.1 | +1.02 | 2.8% |
|  | 0.55 | +8.33 | +0.0053 | 124.5 | +0.65 | 7.9% |
|  | 0.75 | +2.48 | +0.0050 | 40.3 | +0.20 | 8.2% |
|  | 0.95 | +0.21 | −0.0004 | 26.5 | −0.01 | 5.5% |

**Table 5:** Paired sample-0 differences from the four main-grid controls in Table 1: MuseDrift minus each control, *n*=438. Brackets give 95% WT-bootstrap intervals without multiple-comparison correction. Infilling contrasts are in Table 6.

| Metric | Control | 0.3 | 0.4 | 0.55 | 0.75 | 0.95 |
| --- | --- | --- | --- | --- | --- | --- |
| $\Delta\text{pLDDT}$ | ESM-2 650M full-mask | +33.76<br>[32.02, 35.44] | +38.13<br>[36.55, 39.65] | +29.01<br>[27.53, 30.48] | +8.54<br>[7.82, 9.28] | +1.07<br>[0.70, 1.47] |
|  | EvoDiff OA-DM | +26.40<br>[24.75, 28.07] | +26.82<br>[25.27, 28.38] | +21.95<br>[20.48, 23.43] | +10.89<br>[9.92, 11.88] | +1.25<br>[0.82, 1.71] |
|  | BLOSUM62-odds | +35.53<br>[33.90, 37.13] | +35.99<br>[34.50, 37.46] | +24.37<br>[23.01, 25.75] | +9.92<br>[9.15, 10.73] | +1.34<br>[0.99, 1.70] |
|  | Uniform | +36.95<br>[35.28, 38.57] | +41.25<br>[39.75, 42.70] | +36.56<br>[35.08, 38.03] | +14.63<br>[13.71, 15.56] | +1.86<br>[1.50, 2.22] |
| $\Delta\text{TM}$ | ESM-2 650M full-mask | +0.441<br>[0.423, 0.458] | +0.460<br>[0.441, 0.478] | +0.273<br>[0.252, 0.294] | +0.061<br>[0.051, 0.071] | +0.020<br>[0.012, 0.027] |
|  | EvoDiff OA-DM | +0.334<br>[0.314, 0.354] | +0.309<br>[0.290, 0.329] | +0.218<br>[0.198, 0.238] | +0.093<br>[0.079, 0.107] | +0.027<br>[0.019, 0.035] |
|  | BLOSUM62-odds | +0.419<br>[0.400, 0.437] | +0.385<br>[0.365, 0.404] | +0.202<br>[0.183, 0.221] | +0.075<br>[0.063, 0.086] | +0.018<br>[0.011, 0.024] |
|  | Uniform | +0.443<br>[0.425, 0.461] | +0.474<br>[0.457, 0.490] | +0.357<br>[0.337, 0.377] | +0.103<br>[0.091, 0.115] | +0.023<br>[0.016, 0.030] |

## H Selected-site infilling versus WT-prefix/full-mask ESM-2

**Table 6:** CAMEO comparison with ESM-2 selected-site infilling. MuseDrift minus an ESM-2 650M editor with unedited WT residues visible in place. Equal-WT means on 438 WTs at all five targets, using the predesignated first sample. Duplicate records of the same WT are averaged before 20,000 paired WT-bootstrap resamples; brackets give 95% intervals.

| Metric | .30 | .40 | .55 | .75 | .95 |
| --- | --- | --- | --- | --- | --- |
| $\Delta\text{pLDDT}$ | +32.56<br>[+30.96, +34.11] | +36.14<br>[+34.51, +37.79] | +8.33<br>[+7.19, +9.49] | +2.48<br>[+1.93, +3.04] | +0.21<br>[−0.04, +0.46] |
| $\Delta\text{TM}$ | +0.452<br>[+0.433, +0.469] | +0.434<br>[+0.414, +0.454] | +0.084<br>[+0.070, +0.099] | +0.025<br>[+0.018, +0.032] | +0.010<br>[+0.004, +0.017] |

The main ESM-2 control receives a visible WT prefix but starts with the entire variant slot masked and reveals all positions in random order; the edit mask only determines which revealed residues are forced to differ. The sensitivity control masks only those edit sites in an otherwise visible WT sequence, without a separate WT prefix, and predicts them in one forward pass. Both use the same frozen ESM-2 t33 650M snapshot, temperature 1.0, standard-20-AA alphabet, exact Hamming budget and deterministic position seeds. The selected-site infilling pool has *K*=8, but the structural comparison uses only sample index 0 and has no best-of-*K* selection. The two runs use different seed namespaces; their edit-site sets are not paired sample by sample.

## I Budget fidelity per method

At *τ* = .95, 433 of 438 CAMEO WTs have all 64 samples within .05 of the target. Three 38-residue references instead have hit rates of 0, .063, and .234. On the separate K8 extension to *τ* = .97–.995, MAE ranges from .0035 to .0043, but the exact-WT copy rate reaches 2.65% at .995. Aggregate control therefore does not imply a per-WT guarantee or exclude copying arbitrarily close to identity one.

## J Training overlap and homology stratification

### Overlap with actual training sequences

We separately audit the WT and variant strings of all 23,342,731 loader-eligible training pairs. Training strings are stripped of gap characters; complete-string equality is checked exhaustively, followed by MMseqs2 searches at sensitivity 7.5 requiring at least 80% coverage of both query and target. Among the 438 unique CAMEO WTs, 7 match a training WT string exactly and 6 match a training variant string exactly, giving 13 distinct references with an exact match on either side. Combining both sides, 85 references have an exact match or a detected alignment identity of at least 90%; 90 have neither an exact match nor a covered hit detected. Non-exact identities are best-detected similarities under this search protocol, not exhaustive nearest-neighbor maxima. These counts characterize exposure to the training strings; absence of a hit does not establish family-level separation. Table 1 reports the full 438-WT panel; Table 9 reports the filtered subset defined below.

### Evaluation after filtering close training matches

We exclude complete-string matches and detected alignments with identity ≥ 50% to either training side under the coverage criteria above. This retains 226 records representing 223 unique WTs: 133 with detected hits below 50% and 90 with no covered hit. We retain all qualifying WTs and reuse the predesignated first samples for every method in Table 1, averaging duplicate records within WT before equal-WT aggregation and 20,000 paired WT-bootstrap resamples. Table 9 reports structural means, realized identities, and paired structural-score differences on this common subset. Relative to ESM-2 infilling, MuseDrift’s pLDDT gains at *τ* = .55*/.*75 are +10.23 [8.45, 12.09] and +2.85 [1.93, 3.80]. At *τ* = .95, the pLDDT interval includes zero (+.18 [−.23, .59]), while the TM-score difference remains positive (+.014 [.002, .025]). This evaluates sensitivity to detected training similarity rather than generalization to held-out families.

**Table 7:** Budget fidelity on 438 CAMEO wild types, *K*=64. The controls realize the rounded integer edit count by construction and therefore hit target identity to within length-dependent rounding; MuseDrift’s conditioning is learned. This table evaluates the dial as a comparison instrument. Sample MAE is mean(|id(*V,* WT) − *τ* |) over individual samples, not |mean(id) − *τ* |; the two differ sharply when a generator spreads around the target. MAE is not a per-sample upper bound, so the hit-rate rows expose its tails.

| Method | Target identity $\tau$ | | | | |
| --- | --- | --- | --- | --- | --- |
|  | 0.3 | 0.4 | 0.55 | 0.75 | 0.95 |
| *Sample-level identity MAE (lower is better)* |  |  |  |  |  |
| MuseDrift | 0.0208 | 0.0155 | 0.0131 | 0.0111 | 0.0052 |
| ESM-2 650M full-mask | 0.0012 | 0.0012 | 0.0012 | 0.0013 | 0.0013 |
| EvoDiff OA-DM 38M | 0.0012 | 0.0012 | 0.0012 | 0.0013 | 0.0013 |
| BLOSUM62-odds | 0.0012 | 0.0012 | 0.0012 | 0.0013 | 0.0013 |
| Uniform | 0.0012 | 0.0012 | 0.0012 | 0.0013 | 0.0013 |
| *MuseDrift sample hit rate (higher is better)* |  |  |  |  |  |
| Within $\pm 0.02$ | 0.579 | 0.722 | 0.791 | 0.861 | 0.985 |
| Within $\pm 0.05$ | 0.941 | 0.976 | 0.989 | 0.994 | 0.994 |
| *Exact WT copy rate* |  |  |  |  |  |
| MuseDrift | 0.0000 | 0.0000 | 0.0000 | 0.0000 | 0.0000 |
| ESM-2 650M full-mask | 0.0000 | 0.0000 | 0.0000 | 0.0000 | 0.0000 |
| EvoDiff OA-DM 38M | 0.0000 | 0.0000 | 0.0000 | 0.0000 | 0.0000 |
| BLOSUM62-odds | 0.0000 | 0.0000 | 0.0000 | 0.0000 | 0.0000 |
| Uniform | 0.0000 | 0.0000 | 0.0000 | 0.0000 | 0.0000 |

**Table 8:**
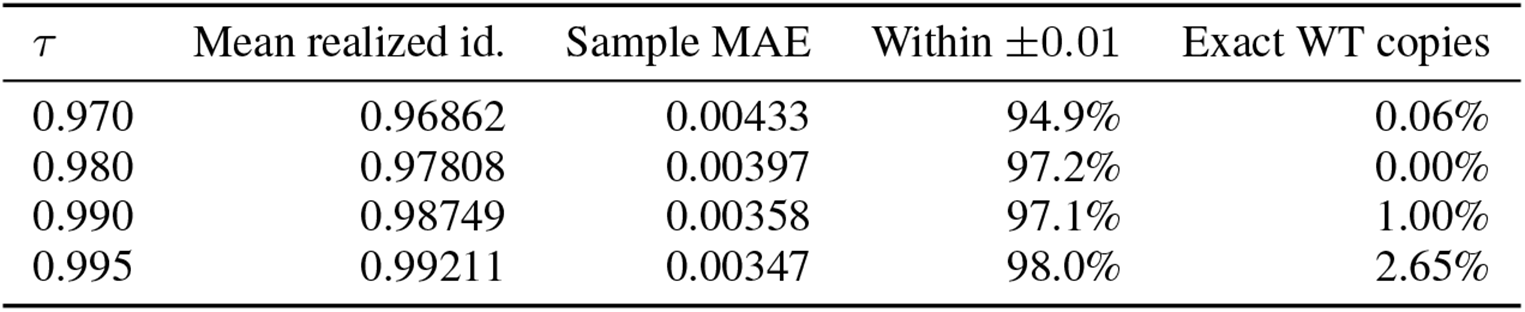
The dial above the grid used for the main comparison: 438 CAMEO wild types, *K*=8, 14,016 sequences. A single substitution on a 250-residue protein is *τ* ≈ 0.996, so this is a shallow regime in which stability campaigns can operate. Sample MAE is tighter here than anywhere in [0.30, 0.95]. The exact-copy rate is the number to watch, since returning the wild type unchanged scores a near-perfect identity for free; it rises to 2.65% at the most demanding target.

**Table 9:** Structural quality after filtering close training matches. Equal-WT means on 223 CAMEO WTs after excluding exact matches and detected alignments with identity ≥ 50% and coverage ≥ 80% on both sequences, against either side of the training pairs. All methods use the same WTs and predesignated first samples. TM-score is relative to the predicted WT structure. Realized identity refers to these scored samples. Bold marks the highest structural-score point estimate, not statistical significance.

| Method | Target identity $\tau$ | | | | |
| --- | --- | --- | --- | --- | --- |
|  | .30 | .40 | .55 | .75 | .95 |
| *ESMFold pLDDT ↑* |  |  |  |  |  |
| MuseDrift | **61.08** | **66.01** | **71.62** | **77.02** | **81.16** |
| ESM-2 infilling | 35.20 | 35.36 | 61.38 | 74.17 | 80.97 |
| ESM-2 full-mask | 35.19 | 34.76 | 45.61 | 67.89 | 80.01 |
| EvoDiff OA-DM | 39.32 | 43.29 | 50.42 | 64.42 | 79.70 |
| BLOSUM62-odds | 32.30 | 34.89 | 47.38 | 65.94 | 79.50 |
| Uniform | 31.54 | 31.33 | 37.87 | 60.99 | 78.92 |
| *Predicted-WT-relative TM-score ↑* |  |  |  |  |  |
| MuseDrift | **0.641** | **0.696** | **0.761** | **0.823** | **0.919** |
| ESM-2 infilling | 0.265 | 0.308 | 0.633 | 0.792 | 0.906 |
| ESM-2 full-mask | 0.271 | 0.299 | 0.485 | 0.748 | 0.895 |
| EvoDiff OA-DM | 0.340 | 0.396 | 0.507 | 0.697 | 0.884 |
| BLOSUM62-odds | 0.282 | 0.344 | 0.532 | 0.725 | 0.892 |
| Uniform | 0.270 | 0.280 | 0.398 | 0.689 | 0.888 |
| *Realized sequence identity* |  |  |  |  |  |
| MuseDrift | 0.3144 | 0.4053 | 0.5548 | 0.7551 | 0.9490 |
| ESM-2 infilling | 0.2998 | 0.3998 | 0.5502 | 0.7498 | 0.9498 |
| ESM-2 full-mask | 0.2998 | 0.3998 | 0.5502 | 0.7498 | 0.9498 |
| EvoDiff OA-DM | 0.2998 | 0.3998 | 0.5502 | 0.7498 | 0.9498 |
| BLOSUM62-odds | 0.2998 | 0.3998 | 0.5502 | 0.7498 | 0.9498 |
| Uniform | 0.2998 | 0.3998 | 0.5502 | 0.7498 | 0.9498 |
| *Internal configuration: WT-token-only* |  |  |  |  |  |
| pLDDT | 40.37 | 48.68 | 57.47 | 69.96 | 79.77 |
| TM-score | 0.419 | 0.515 | 0.627 | 0.767 | 0.913 |
| Realized identity | 0.2817 | 0.3690 | 0.4730 | 0.6653 | 0.9034 |
| *MuseDrift minus ESM-2 infilling [95% paired interval]* |  |  |  |  |  |
| ΔpLDDT | +25.88 | +30.65 | +10.23 | +2.85 | +0.18 |
|  | [23.65, 28.09] | [28.38, 32.90] | [8.45, 12.09] | [1.93, 3.80] | [−0.23, 0.59] |
| ΔTM | +0.376 | +0.388 | +0.128 | +0.031 | +0.014 |
|  | [0.349, 0.402] | [0.361, 0.415] | [0.104, 0.153] | [0.019, 0.044] | [0.002, 0.025] |

WT-token-only is a separately trained, temperature-calibrated configuration; training schedules and realized identities differ from the full model (Appendix Q). Intervals use 20,000 paired WT-bootstrap resamples after averaging duplicate records within WT. Filtering concerns detected sequence similarity, not family membership.

### Raw-seed homology stratification

All 438 evaluation wild types were searched against the 1,330,187-entry raw pre-cutoff UniRef50 seed universe with MMseqs2 at sensitivity 7, retaining the best hit at ≥50% query coverage. This database is upstream of the train/val/test split and includes seeds that were not eligible for training, so the result must not be interpreted as distance from the training set. Bin sizes: no hit 12, *<*30% 9, 30–50% 53, 50–80% 166, ≥80% 198.

The two most distant bins contain only 21 WTs in total and are not bootstrapped separately. The no-hit bin is also length-skewed: its median length is 142 residues, compared with 266 for the full panel. Neither no-hit status nor this stratification establishes sequence novelty.

**Table 10:**
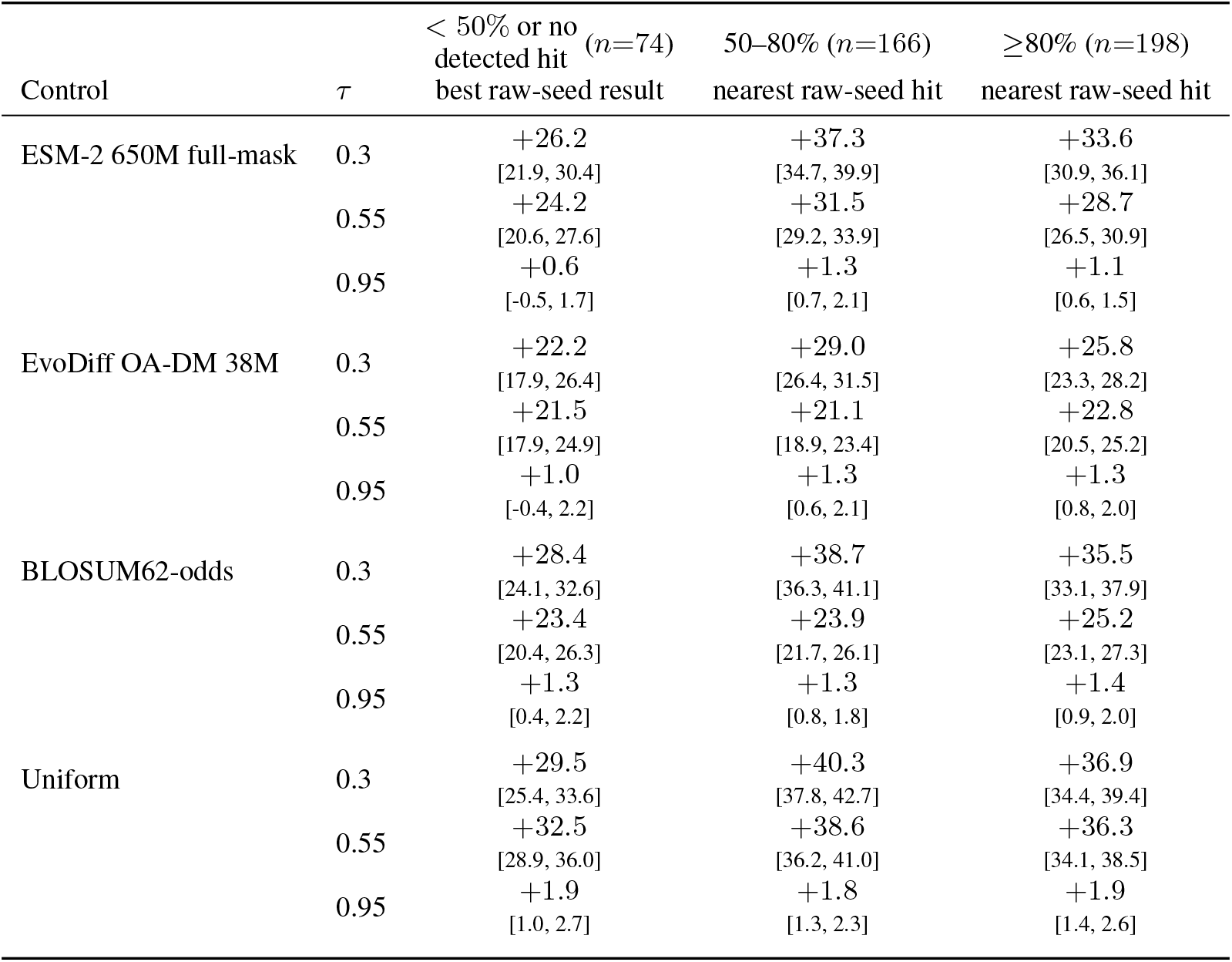
The main comparison stratified by how close each evaluation wild type sits to the raw pre-cutoff UniRef50 seed universe (MMseqs2 best hit, ≥50% query coverage). Entries use sample index 0 and are ΔpLDDT for MuseDrift minus the control, paired within wild type, 95% bootstrap intervals over the wild types in that bin. At deep drift the advantage remains positive in the leftmost column but is often smaller; at *τ* =0.95 two learned-control intervals include zero. The database is broader than the eligible training split; this is a raw-seed-similarity sensitivity, not a training-exposure audit or family holdout.

| Control | $\tau$ | < 50% or no<br>detected hit<br>($n=74$) | 50–80% ($n=166$) | $\geq 80\%$ ($n=198$) |
| --- | --- | --- | --- | --- |
|  |  | best raw-seed result | nearest raw-seed hit | nearest raw-seed hit |
| ESM-2 650M full-mask | 0.3 | +26.2 | +37.3 | +33.6 |
|  |  | [21.9, 30.4] | [34.7, 39.9] | [30.9, 36.1] |
|  | 0.55 | +24.2 | +31.5 | +28.7 |
|  |  | [20.6, 27.6] | [29.2, 33.9] | [26.5, 30.9] |
|  | 0.95 | +0.6 | +1.3 | +1.1 |
|  |  | [-0.5, 1.7] | [0.7, 2.1] | [0.6, 1.5] |
| EvoDiff OA-DM 38M | 0.3 | +22.2 | +29.0 | +25.8 |
|  |  | [17.9, 26.4] | [26.4, 31.5] | [23.3, 28.2] |
|  | 0.55 | +21.5 | +21.1 | +22.8 |
|  |  | [17.9, 24.9] | [18.9, 23.4] | [20.5, 25.2] |
|  | 0.95 | +1.0 | +1.3 | +1.3 |
|  |  | [-0.4, 2.2] | [0.6, 2.1] | [0.8, 2.0] |
| BLOSUM62-odds | 0.3 | +28.4 | +38.7 | +35.5 |
|  |  | [24.1, 32.6] | [36.3, 41.1] | [33.1, 37.9] |
|  | 0.55 | +23.4 | +23.9 | +25.2 |
|  |  | [20.4, 26.3] | [21.7, 26.1] | [23.1, 27.3] |
|  | 0.95 | +1.3 | +1.3 | +1.4 |
|  |  | [0.4, 2.2] | [0.8, 1.8] | [0.9, 2.0] |
| Uniform | 0.3 | +29.5 | +40.3 | +36.9 |
|  |  | [25.4, 33.6] | [37.8, 42.7] | [34.4, 39.4] |
|  | 0.55 | +32.5 | +38.6 | +36.3 |
|  |  | [28.9, 36.0] | [36.2, 41.0] | [34.1, 38.5] |
|  | 0.95 | +1.9 | +1.8 | +1.9 |
|  |  | [1.0, 2.7] | [1.3, 2.3] | [1.4, 2.6] |

## K Within-library diversity

Every main-grid reference-record–target library contains 64 distinct sequences and no exact WT copies. Paired uncertainty is available for the library statistics: Table 12 gives the *τ* = .55 contrasts. Under these generation protocols, MuseDrift has higher pairwise identity and lower edit-set Jaccard distance than each random-site control. This compares the complete generation procedures; it does not isolate the effect of site selection or establish a causal trade-off with structural quality.

## L Auxiliary sequence diagnostics

These auxiliary sequence metrics include control-favoring comparisons; the structural results do not imply an advantage on every sequence score.

**Table 11:** Diversity within each library. , 438 CAMEO wild types, *K*=64, wild-type macro-means. MuseDrift has higher pairwise identity and lower cluster-count and edit-set-Jaccard point estimates than every control; all three directions indicate lower diversity. Paired WT-bootstrap intervals for *τ* = .55 are reported in Table 12. These comparisons do not establish a causal quality–diversity trade-off. Pairwise identity is exact token agreement over all wild-type columns. Clusters are single-linkage connected components over all ^64^ pairs at an identity threshold equal to the target *τ* ; a fixed 0.90 threshold saturates at 1.000 for every method at *τ* ≤ 0.75 and discriminates nothing. Because the reported threshold changes with *τ*, cluster-ratio deficits are not ranked across targets and threshold robustness is unresolved. Jaccard distance compares which positions changed, ignoring what they became.

| Method | 0.3 | 0.4 | 0.55 | 0.75 | 0.95 |
| --- | --- | --- | --- | --- | --- |
| *Mean pairwise identity* (lower is more diverse) |  |  |  |  |  |
| MuseDrift | 0.2305 | 0.2968 | 0.4181 | 0.6300 | 0.9082 |
| ESM-2 650M full-mask | 0.1227 | 0.1834 | 0.3167 | 0.5686 | 0.9025 |
| EvoDiff OA-DM | 0.1271 | 0.1906 | 0.3231 | 0.5700 | 0.9025 |
| BLOSUM62-odds | 0.1249 | 0.1856 | 0.3171 | 0.5668 | 0.9023 |
| Uniform | 0.1158 | 0.1789 | 0.3133 | 0.5656 | 0.9023 |
| *Cluster count / K* (higher is more diverse) |  |  |  |  |  |
| MuseDrift | 0.6321 | 0.8464 | 0.9601 | 0.9870 | 0.9784 |
| ESM-2 650M full-mask | 0.9998 | 1.0000 | 1.0000 | 1.0000 | 0.9972 |
| EvoDiff OA-DM | 0.9988 | 1.0000 | 1.0000 | 1.0000 | 0.9978 |
| BLOSUM62-odds | 1.0000 | 1.0000 | 1.0000 | 1.0000 | 0.9984 |
| Uniform | 1.0000 | 1.0000 | 1.0000 | 1.0000 | 0.9981 |
| *Edit-set Jaccard distance* (higher is more diverse) |  |  |  |  |  |
| MuseDrift | 0.3782 | 0.4562 | 0.5796 | 0.7438 | 0.9283 |
| ESM-2 650M full-mask | 0.4610 | 0.5707 | 0.7089 | 0.8559 | 0.9729 |
| EvoDiff OA-DM | 0.4610 | 0.5707 | 0.7089 | 0.8559 | 0.9729 |
| BLOSUM62-odds | 0.4610 | 0.5707 | 0.7089 | 0.8559 | 0.9729 |
| Uniform | 0.4610 | 0.5707 | 0.7089 | 0.8559 | 0.9729 |

**Table 12:** Paired library-diversity differences at. *τ* = .55. Values are MuseDrift minus each control on 438 WTs, *K* = 64, with 95% intervals from 20,000 paired WT bootstrap replicates. Positive pairwise-identity differences and negative edit-set-distance differences both indicate lower diversity for MuseDrift under these generation protocols.

| Control | $\Delta$ pairwise identity | $\Delta$ edit-set Jaccard distance |
| --- | --- | --- |
| Uniform | +0.1048<br>[+0.1013, +0.1084] | -0.1292<br>[-0.1343, -0.1243] |
| BLOSUM62-odds | +0.1010<br>[+0.0975, +0.1046] | -0.1292<br>[-0.1343, -0.1243] |
| ESM-2 full-mask | +0.1014<br>[+0.0978, +0.1050] | -0.1292<br>[-0.1343, -0.1243] |
| EvoDiff OA-DM | +0.0950<br>[+0.0916, +0.0985] | -0.1292<br>[-0.1343, -0.1243] |

**Table 13:**
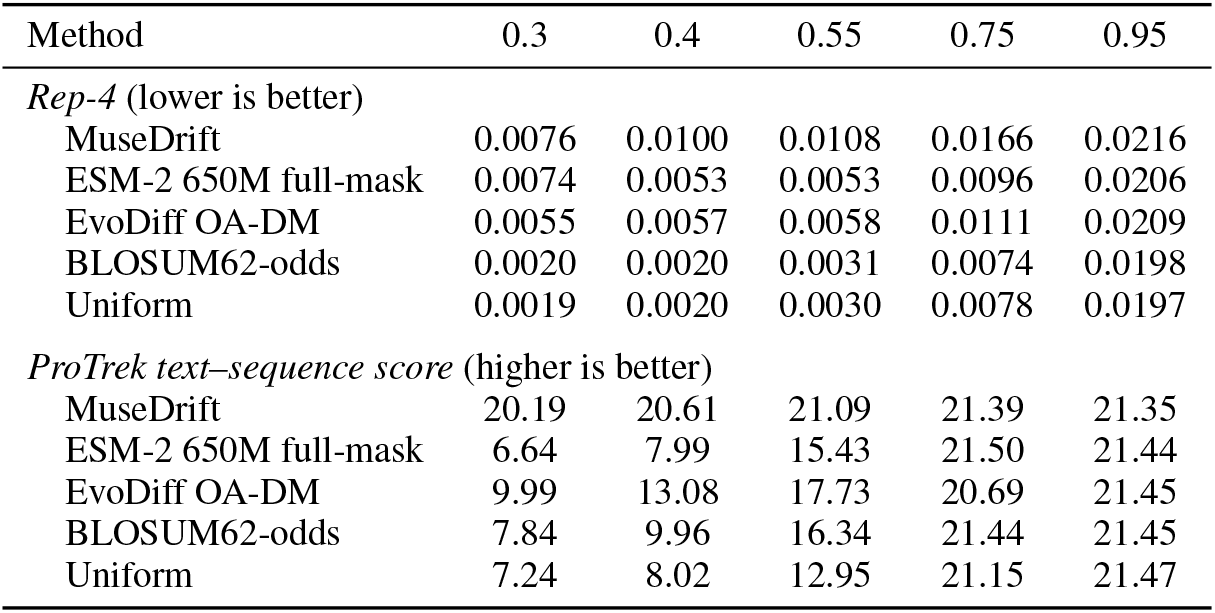
Auxiliary sequence diagnostics for the predesignated sample-index-0 members. , 438 WT hashes. Rep-4 is the fraction of 4-mer windows whose type repeats within the sequence (lower is better). ProTrek (Su et al., 2026) is 100 times text–sequence embedding cosine similarity (higher is better) to the released function string. At *τ* =0.55, all four paired Rep-4 intervals favour the controls. At *τ* =0.95, only the BLOSUM62 ProTrek interval excludes zero, in the control-favouring direction; the other three include zero. Neither metric is experimental evidence of function.

| Method | 0.3 | 0.4 | 0.55 | 0.75 | 0.95 |
| --- | --- | --- | --- | --- | --- |
| *Rep-4* (lower is better) |  |  |  |  |  |
| MuseDrift | 0.0076 | 0.0100 | 0.0108 | 0.0166 | 0.0216 |
| ESM-2 650M full-mask | 0.0074 | 0.0053 | 0.0053 | 0.0096 | 0.0206 |
| EvoDiff OA-DM | 0.0055 | 0.0057 | 0.0058 | 0.0111 | 0.0209 |
| BLOSUM62-odds | 0.0020 | 0.0020 | 0.0031 | 0.0074 | 0.0198 |
| Uniform | 0.0019 | 0.0020 | 0.0030 | 0.0078 | 0.0197 |
| *ProTrek text–sequence score* (higher is better) |  |  |  |  |  |
| MuseDrift | 20.19 | 20.61 | 21.09 | 21.39 | 21.35 |
| ESM-2 650M full-mask | 6.64 | 7.99 | 15.43 | 21.50 | 21.44 |
| EvoDiff OA-DM | 9.99 | 13.08 | 17.73 | 20.69 | 21.45 |
| BLOSUM62-odds | 7.84 | 9.96 | 16.34 | 21.44 | 21.45 |
| Uniform | 7.24 | 8.02 | 12.95 | 21.15 | 21.47 |

**Figure 4:**
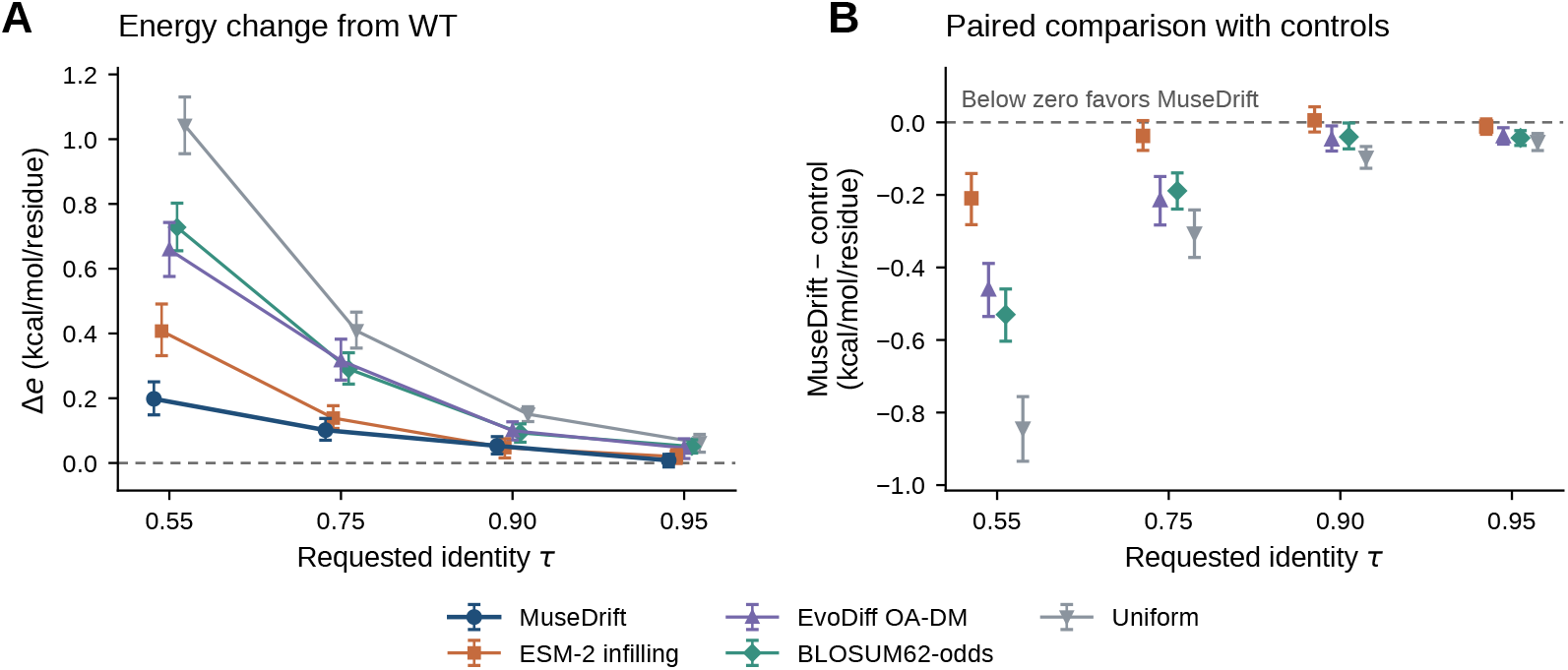
FoldX energy assessment on predicted structures. The same 80 CAMEO WTs are evaluated at four target identities, using predesignated sample 0 for each method and target. **A:** Mean length-normalized energy change from the predicted WT, Δ*e* = [*E*(*V*) − *E*(*W*)]*/L_W_* ; lower values indicate lower FoldX energy. The dashed line marks the WT reference (Δ*e* = 0). **B:** Paired differences, Δ*e*_MuseDrift_ −Δ*e*_control_; values below zero favor MuseDrift. Colors identify the same controls as in A. Horizontal offsets separate methods at each target in both panels. Error bars in both panels are 95% WT-bootstrap intervals from 10,000 resamples. Structures are predicted with ESMFold; these scores provide a computational energy assessment, not measured stability. Full paired contrasts are in Table 14.

**Table 14:** FoldX-reported energy change on predicted ESMFold structures. For a separate 80-WT, four-target sequence draw, entries are MuseDrift minus each control in [*E*(*V*) − *E*(WT)]*/L*, paired within wild type, 10,000 bootstrap replicates over 80 wild-type hashes, sample index 0. **Negative favours MuseDrift**. FoldX is not a language-model scorer, but its input structures are ESM-Fold predictions; this is therefore a second computational proxy, not experimental stability. The result agrees against Uniform, BLOSUM62, and EvoDiff, but the intervals against ESM-2 infilling contain zero at *τ* ≥ 0.75.

| Control | 0.55 | 0.75 | 0.90 | 0.95 |
| --- | --- | --- | --- | --- |
| ESM-2 650M infilling | -0.209<br>[-0.282, -0.141] | -0.037<br>[-0.077, +0.005] | +0.006<br>[-0.027, +0.043] | -0.011<br>[-0.033, +0.009] |
| EvoDiff OA-DM | -0.461<br>[-0.535, -0.389] | -0.215<br>[-0.283, -0.149] | -0.047<br>[-0.079, -0.010] | -0.038<br>[-0.060, -0.015] |
| BLOSUM62-odds | -0.530<br>[-0.603, -0.459] | -0.189<br>[-0.239, -0.139] | -0.040<br>[-0.073, -0.002] | -0.043<br>[-0.064, -0.023] |
| Uniform | -0.844<br>[-0.934, -0.756] | -0.307<br>[-0.373, -0.242] | -0.098<br>[-0.127, -0.067] | -0.054<br>[-0.078, -0.031] |

## M FoldX energy sensitivity on a separate draw

Figure 4 visualizes the WT-relative energy changes and their paired differences from the four controls on the same complete 80-WT panel. Both panels use sample index 0; no energy-based selection is applied.

The frozen FoldX 5.1 binary (Schymkowitz et al., 2005) (SHA-256 prefix `3f892047bc23`) was run independently for each structure in an isolated directory using `RepairPDB` followed by `Stability` on the repaired structure, with `--screen=false`, `LANG=C`, and a 600-second time-out; raw failures were retained. The reported quantity is [*E*_FoldX_(*V*) − *E*_FoldX_(WT)]*/L*_WT_, not a measured folding free energy.

## N Substitution patterns and MSA conservation

### Substitution-matrix statistic

The primary panel contains 441 records corresponding to all 438 unique CAMEO WT hashes, with eight samples per target *τ* ∈ {.55, .75, .90, .95}. For each WT, we count aligned WT-to-output residue pairs, including diagonal observations. After equal-WT aggregation, the matrix is converted to row-conditional probabilities and log_2_ odds against the generated-residue marginal. Zero-count cells map to zero, values are clipped to [−5, 5], and the matrix is averaged with its transpose. Pearson correlation with BLOSUM62 is computed over observed unordered off-diagonal pairs.

Uncertainty uses 20,000 Rubin Bayesian-bootstrap draws over WT hashes, with the observed-pair support fixed to that of the point estimate. These equal-tail intervals describe a nonlinear pooled statistic and need not contain its equal-weight plug-in estimate. The alternative statistic symmetrizes the joint count distribution and uses all 400 cells; including self-matches and composition makes even Uniform strongly positive. We report it separately rather than changing definitions between methods. A previously fixed 408-WT subset excluding the 30 CAMEO references used for calibrating configurations without FiLM (Appendix Q) gives MuseDrift correlations of .918/.861/.689/.582 at the four targets. Across methods and targets, the absolute difference from the full-panel estimate is at most .0101.

### MSA entropy and mutation placement

On the 436 eligible CAMEO WTs, we compute Shannon entropy from each MSA column’s 20-residue count vector. Query residues, gaps, and nonstandard amino acids do not contribute to these counts. For each WT and target, mutation frequency is the fraction of eight generated variants differing from the WT at that position; duplicate record aliases are averaged first. Spearman correlation is computed within WT over valid columns, then summarized across WTs. All 436 correlations are defined at every target. For the entropy-binned curve, frequencies are averaged within each WT and bin, then equally across WTs represented in the bin. Intervals use 20,000 WT-bootstrap resamples with fixed generated sequences and MSA profiles.

**Table 15:** BLOSUM62 substitution-pattern correlations. All 438 CAMEO WTs, *K* = 8 at each target. Top: row-conditional, symmetrized log-odds Pearson correlation over observed unordered off-diagonal pairs, with 95% equal-tail intervals from 20,000 WT-hash Bayesian-bootstrap draws. Bottom: the alternative symmetric-joint 400-cell statistic on the same 438 WTs, point estimates only. The two definitions are not interchangeable.

| Method | .55 | .75 | .90 | .95 |
| --- | --- | --- | --- | --- |
| MuseDrift | 0.919<br>[0.911, 0.922] | 0.863<br>[0.852, 0.870] | 0.693<br>[0.680, 0.705] | 0.592<br>[0.584, 0.601] |
| ESM-2 infilling | 0.863<br>[0.854, 0.867] | 0.875<br>[0.866, 0.880] | 0.874<br>[0.861, 0.879] | 0.833<br>[0.817, 0.841] |
| EvoDiff OA-DM | 0.858<br>[0.835, 0.863] | 0.872<br>[0.850, 0.876] | 0.866<br>[0.832, 0.869] | 0.871<br>[0.821, 0.868] |
| BLOSUM62-odds | 0.871<br>[0.864, 0.875] | 0.830<br>[0.819, 0.837] | 0.782<br>[0.766, 0.793] | 0.740<br>[0.716, 0.751] |
| Uniform | -0.087<br>[-0.108, -0.065] | -0.112<br>[-0.131, -0.089] | -0.096<br>[-0.120, -0.069] | -0.122<br>[-0.150, -0.093] |
| *Alternative 400-cell statistic* |  |  |  |  |
| MuseDrift | 0.697 | 0.657 | 0.628 | 0.618 |
| ESM-2 infilling | 0.664 | 0.641 | 0.620 | 0.614 |
| EvoDiff OA-DM | 0.630 | 0.621 | 0.613 | 0.611 |
| BLOSUM62-odds | 0.666 | 0.632 | 0.616 | 0.612 |
| Uniform | 0.604 | 0.606 | 0.608 | 0.608 |

The random-placement reference is the expected frequency when each variant’s actual edit count is distributed uniformly over the WT. Within a WT this expectation is constant across positions; the plotted curve uses the same WT and bin weights as the observed frequencies. It therefore controls mutation burden without imposing the requested count.

MSA entropy also depends on alignment composition and redundancy. The association describes where mutations occur relative to sequence conservation; it does not test causal evolutionary mechanisms or experimental function.

**Table 16:**
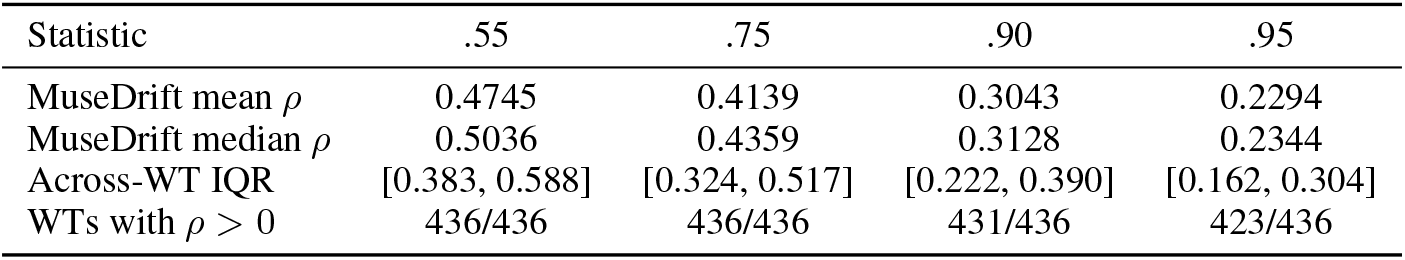
Mutation placement versus MSA entropy on CAMEO. Of 438 unique WTs, 436 meet the MSA coverage criteria; two have no valid entropy columns. Within-WT Spearman correlations use all eight generated variants at each target. IQR describes variation across WTs.

| Statistic | .55 | .75 | .90 | .95 |
| --- | --- | --- | --- | --- |
| MuseDrift mean $\rho$ | 0.4745 | 0.4139 | 0.3043 | 0.2294 |
| MuseDrift median $\rho$ | 0.5036 | 0.4359 | 0.3128 | 0.2344 |
| Across-WT IQR | [0.383, 0.588] | [0.324, 0.517] | [0.222, 0.390] | [0.162, 0.304] |
| WTs with $\rho > 0$ | 436/436 | 436/436 | 431/436 | 423/436 |

## O External fitness prediction on generated libraries

### Panel and sample scope

This analysis uses a fixed 429-WT subset of the 438-WT CAMEO panel. Before scoring, nine references flagged by virus- or toxin-related domain annotations were provisionally excluded pending source-organism verification: CAMEO record IDs 00008, 00072, 00144, 00229, 00278, 00282, 00359, 00360, and 00494. This subset also defines the additional ProDVa evaluation (Appendix R); the WT-conditioned structural and substitution comparisons retain all 438 WTs. The retained panel comprises 432 reference records, including three duplicate-sequence pairs. All six methods use the same references and five targets *T_C_*. All existing draws are scored: *K* = 64 for each main-grid method and *K* = 8 for ESM-2 infilling, totaling 708,480 draws across reference records. Duplicate sequences remain in their libraries; there is no score-based selection.

### Predictors and score definition

ProSST (Li et al., 2024) receives the unmasked WT sequence and quantized tokens from its ESMFold-predicted structure. The same WT structure is reused for every generator; variants are not individually folded for this score. The suffixes 2048 and 4096 denote structural vocabulary sizes, not numbers of generated samples. Following the released scoring code, let *ℓ_r,i_*(*a*) be the predictor’s WT-context log score for residue *a* at position *i*. For a generated sequence *V*, we compute

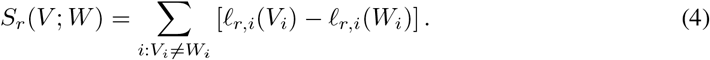

For ProSST, *ℓ* is the log-softmax of its output logits from a single unmasked WT forward pass. VenusREM (Tan et al., 2025) uses *ℓ*_Venus_ = 0.2*ℓ*_ProSST-2048_ + 0.8*ℓ*_MSA_, where *ℓ*_MSA_ is the log-softmax of aligned residue frequencies, as in the released implementation. We reuse the full-query ColabFold A3M alignments, remove insertions, and retain the query. Gaps and nonstandard residues each occupy one alignment column and are encoded as padding and unknown tokens, respectively. This column-preserving projection is distinct from the gap-excluding entropy analysis. Implementation checks against the released scoring functions agree exactly for ProSST and within 1.91 × 10^−6^ for VenusREM on 30 prespecified variants spanning all methods and targets.

### Aggregation and uncertainty

We average samples within each reference record, then records within each exact-WT sequence hash, and finally the 429 WT means with equal weight. Thus the three duplicate-reference pairs receive no extra weight. All targets and predictors are reported separately; their score scales are not directly comparable. Negative scores closer to zero are more favorable relative to the WT, not measurements of absolute fitness. Paired 95% intervals use 20,000 resamples of the 429 WT hashes (seed 20260924), holding models and generated sequences fixed. Intervals are unadjusted for multiple comparisons and do not model dependence between homologous WTs.

**Table 17:** ProSST-4096 sensitivity on the same 429-WT subset. All-sample, WT-relative additive scores with the same aggregation and sample scopes as Table 2. Higher is better; bold marks the highest mean.

| Generator | Target identity $\tau$ | | | | |
| --- | --- | --- | --- | --- | --- |
|  | .30 | .40 | .55 | .75 | .95 |
| *ProSST-4096* ↑ | | | | | |
| MuseDrift | **−435.93** | **−356.71** | **−250.50** | **−127.98** | **−24.51** |
| ESM-2 infilling | −757.61 | −613.50 | −330.64 | −166.57 | −32.16 |
| ESM-2 full-mask | −735.01 | −622.07 | −440.99 | −216.24 | −40.01 |
| EvoDiff OA-DM | −666.84 | −549.68 | −392.09 | −207.30 | −39.89 |
| BLOSUM62-odds | −662.77 | −568.78 | −426.30 | −236.68 | −47.47 |
| Uniform | −751.23 | −644.12 | −482.75 | −268.29 | −53.87 |

**Table 18:**
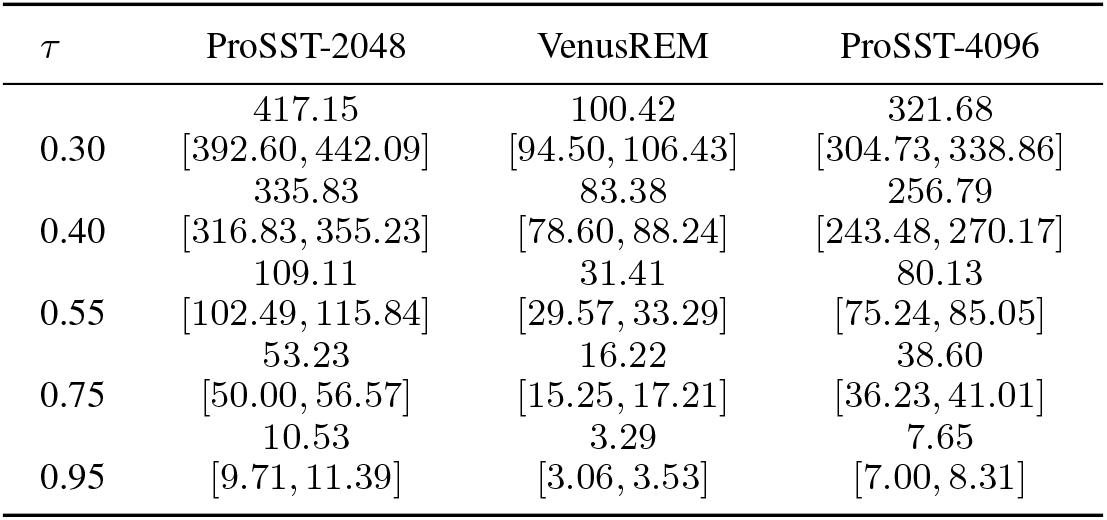
External-score advantages over ESM-2 infilling. Paired differences (MuseDrift minus ESM-2 infilling) in all-sample mean score, with individual 95% intervals from 20,000 WT-bootstrap resamples. All 429 WTs are paired; intervals are not adjusted for multiple comparisons.

| $\tau$ | ProSST-2048 | VenusREM | ProSST-4096 |
| --- | --- | --- | --- |
|  | 417.15 | 100.42 | 321.68 |
| 0.30 | [392.60, 442.09] | [94.50, 106.43] | [304.73, 338.86] |
|  | 335.83 | 83.38 | 256.79 |
| 0.40 | [316.83, 355.23] | [78.60, 88.24] | [243.48, 270.17] |
|  | 109.11 | 31.41 | 80.13 |
| 0.55 | [102.49, 115.84] | [29.57, 33.29] | [75.24, 85.05] |
|  | 53.23 | 16.22 | 38.60 |
| 0.75 | [50.00, 56.57] | [15.25, 17.21] | [36.23, 41.01] |
|  | 10.53 | 3.29 | 7.65 |
| 0.95 | [9.71, 11.39] | [3.06, 3.53] | [7.00, 8.31] |

### Sample-count and mutation-burden sensitivity

We repeat the comparison using only sample indices 0–7 for every method. Separately, we divide each variant’s score by its actual number of substitutions before averaging. Across all three predictors and five targets, MuseDrift retains a higher mean than each of the five baselines under both adjustments and their combination; every individual paired 95% interval remains above zero. Table 19 reports these comparisons against ESM-2 infilling. ProSST-4096 results are consistent with the primary scoring configuration (Table 17). VenusREM shares the ProSST-2048 backbone, so these configurations are not independent model families. All scores sum single-site effects in WT context and do not capture epistasis; this limits their interpretation for distant, multi-substitution variants in particular. No generated library has been experimentally assayed for retained function.

**Table 19:** Sample-count and mutation-burden sensitivity. Paired mean differences from ESM-2 infilling, with individual 95% WT-bootstrap intervals. Common *K* = 8 retains sample indices 0–7 from each method. Per-edit scores divide each variant score by its actual substitution count before aggregation. Positive differences favor MuseDrift. All comparisons use 429 WTs.

| $\tau$ | Common $K = 8$, raw | All samples, per edit | Common $K = 8$, per edit |
| --- | --- | --- | --- |
| *ProSST-2048* |  |  |  |
|  | 417.75 | 2.084 | 2.089 |
| 0.30 | [393.18, 442.70] | [2.021, 2.148] | [2.025, 2.153] |
|  | 336.80 | 1.998 | 2.001 |
| 0.40 | [317.68, 356.25] | [1.941, 2.055] | [1.943, 2.058] |
|  | 108.57 | 0.891 | 0.886 |
| 0.55 | [101.89, 115.33] | [0.853, 0.928] | [0.848, 0.923] |
|  | 53.35 | 0.722 | 0.717 |
| 0.75 | [50.12, 56.70] | [0.686, 0.758] | [0.681, 0.752] |
|  | 10.58 | 0.745 | 0.745 |
| 0.95 | [9.70, 11.49] | [0.695, 0.795] | [0.690, 0.799] |
| *VenusREM* |  |  |  |
|  | 100.56 | 0.503 | 0.504 |
| 0.30 | [94.64, 106.60] | [0.487, 0.519] | [0.488, 0.520] |
|  | 83.63 | 0.496 | 0.497 |
| 0.40 | [78.83, 88.49] | [0.481, 0.511] | [0.482, 0.512] |
|  | 31.26 | 0.254 | 0.253 |
| 0.55 | [29.41, 33.15] | [0.245, 0.264] | [0.243, 0.263] |
|  | 16.23 | 0.223 | 0.222 |
| 0.75 | [15.26, 17.23] | [0.213, 0.232] | [0.212, 0.231] |
|  | 3.30 | 0.233 | 0.233 |
| 0.95 | [3.06, 3.55] | [0.220, 0.247] | [0.218, 0.247] |
| *ProSST-4096* |  |  |  |
|  | 321.79 | 1.691 | 1.692 |
| 0.30 | [304.71, 338.97] | [1.642, 1.741] | [1.642, 1.743] |
|  | 257.48 | 1.599 | 1.602 |
| 0.40 | [244.08, 270.88] | [1.552, 1.646] | [1.554, 1.650] |
|  | 79.64 | 0.703 | 0.700 |
| 0.55 | [74.63, 84.68] | [0.666, 0.741] | [0.662, 0.739] |
|  | 38.71 | 0.569 | 0.567 |
| 0.75 | [36.33, 41.13] | [0.534, 0.606] | [0.530, 0.605] |
|  | 7.73 | 0.572 | 0.576 |
| 0.95 | [7.03, 8.45] | [0.524, 0.620] | [0.525, 0.627] |

## P Zero-shot mutation-effect prediction on ProteinGym

ProteinGym evaluates whether MuseDrift’s own conditional scores associate with measured mutation effects on independently assayed variants (Notin et al., 2023). This tests the generator as a zero-shot scorer, separately from external evaluation of its generated libraries.

Mean within-assay Spearman correlations are .2677 [.2473, .2881] for single mutants and .1846 [.1269, .2417] for multiple mutants (Table 20). ESM-1v and ESM-2 have higher point estimates; the multi-mutant difference from ESM-1v has an interval spanning zero. The positive association therefore does not establish MuseDrift as a competitive standalone mutation-effect predictor.

**Table 20:**
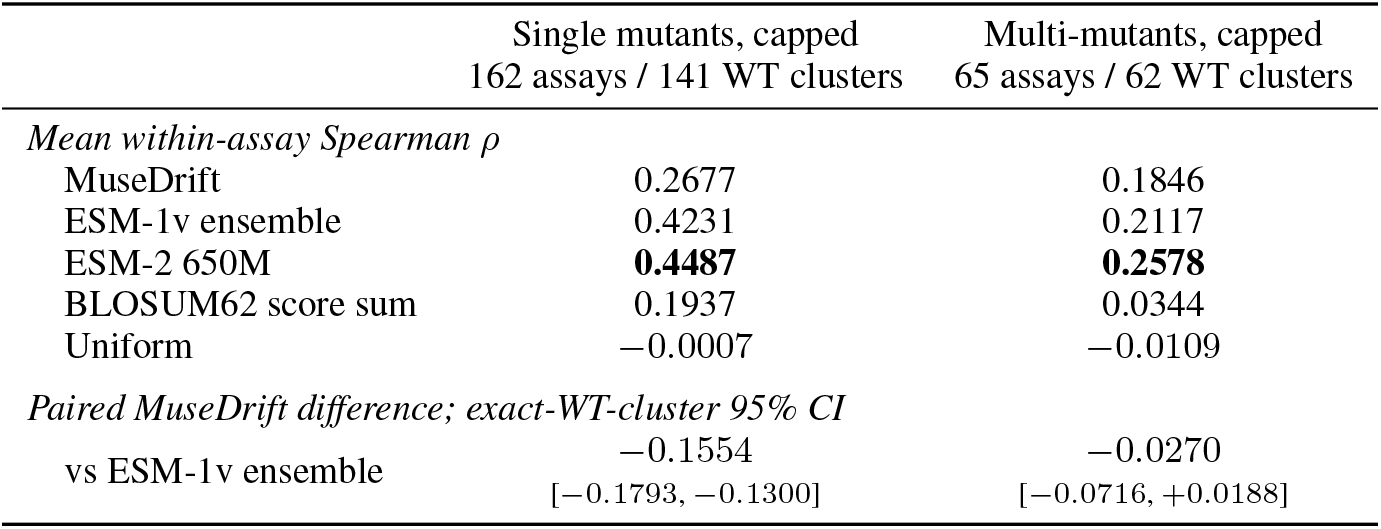
Zero-shot score–fitness association on ProteinGym. Spearman correlations are computed within assays and then macro-averaged. Each assay contributes at most 5,000 uniformly sampled variants. The multi-mutant column uses the WT-marginal score on 2–8 substitutions. The paired contrast resamples exact-WT clusters for its 95% interval.

For an assayed variant *V*, let *M* = {*i* : *V_i_* ≠ *W_i_*} and let *C_M_* (*W*) equal the WT outside *M*, with all focal sites masked. At *τ* = 1 − |*M* |*/L*, the WT-marginal score is

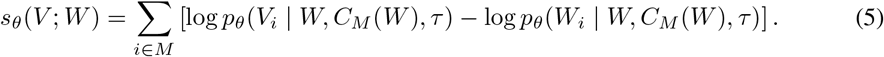

All substituted sites are masked simultaneously, and no assay labels enter the score.

Eligibility requires a strict-20-AA reference of length *L* ≤ 511, a finite DMS score, parsable one-indexed substitutions whose declared WT residues match the reference, and at least 20 retained variants per assay. Panel A keeps exactly one substitution; Panel B keeps 2 ≤ *m* ≤ 8 and rejects repeated or conflicting sites.

For the primary WT-marginal score, every focal substitution is masked simultaneously in an otherwise WT-context variant slot. ESM-2 t33 uses the same simultaneous-focal-mask mutation log-odds definition. ESM-1v is the released ProteinGym ensemble score (Meier et al., 2021); BLOSUM62 (Henikoff & Henikoff, 1992) is the sum of matrix substitution scores; and Uniform is a deterministic hash-seeded random null for each assay–mutant key, not the structural benchmark’s random mutator. Alongside the identity condition, MuseDrift receives only the WT sequence and its frozen ESM-2 t12 features as reference context.

Both panels traverse the frozen 217-assay index in its stored order and share one `numpy.random.default rng(seed=0)` stream. After eligibility filtering, each assay is uniformly subsampled to at most 5,000 rows. Panel A therefore retains 328,027 of 383,569 eligible single mutants (27 assays capped; 55,542 rows excluded), while Panel B retains 123,541 of 1,510,840 eligible 2–8-mutants (14 assays capped; 1,387,299 rows excluded). Exact reproduction requires both the frozen index order and seed. Every Panel-B assay also appears in Panel A. Intervals resample exact-WT sequence clusters and retain every assay belonging to a sampled cluster. The mutant-context sensitivity evaluates each focal mutation while exposing the other substitutions, but its lower point estimate supplies no evidence of biological epistasis.

Stability-labeled assays do not explain the entire association: MuseDrift reaches *ρ* = 0.2949 [0.2664, 0.3231] on 97 other assays spanning 76 WT clusters. This non-Stability group comprises Activity, Binding, Expression, and OrganismalFitness assays. Within its 31 Activity assays over 30 WT clusters, *ρ* = 0.3053 [0.2605, 0.3505]. Both analyses use the single-mutant panel. The multimutant non-Stability group contains 15 assays over 12 exact-WT clusters and yields *ρ* = 0.3454 [0.2494, 0.4359]. Its Activity-only subset has just five assays; we do not use that small subset as a primary estimate. In a separate target-string sensitivity, 26 Panel-A assays have a ProteinGym target sequence matching at least one eligible training-side alignment string exactly; removing them leaves *ρ* = 0.2610 [0.2382, 0.2837] across 136 assays and 120 clusters. This audit does not inspect every assayed mutant string or establish family separation. Likewise, MuseDrift’s smaller point gap to ESM-1v on the capped multi-mutant panel cannot be attributed causally to pair training: the panels have different variant distributions and no retraining intervention was performed. The EvoDiff generation comparison does not supply a validated multi-mutant likelihood scorer for this table.

## Q Comparison of conditioning configurations

We compare four configurations of the same 12-layer, 768-wide backbone: the WT-token-only configuration; WT tokens with numerical identity FiLM; WT tokens with ESM-2 WT features; and the full model with all three inputs. Table 21 identifies which inputs each configuration receives. Removing ESM-2 embeddings removes the WT-feature adapter as well. Both configurations with ESM-2 embeddings use the same feature assignment (Appendix C). All configurations retain the WT token context and use 50-step random-order decoding over the 20 standard amino acids. Models with FiLM receive numerical target identity directly; models without FiLM use a calibrated identity-to-temperature mapping.

The structural comparison uses the same 438 unique CAMEO WTs and predesignated first sample as Table 1. Duplicate reference records are averaged within each WT before averaging across WTs. Structural scores for all four configurations are complete at nominal identity .55, so the four-way comparison is restricted to this common setting (Table 21); the full model and the WT-token-only configuration additionally cover all five settings in Table 1. Table 22 reports their realized identities.

**Table 21:**
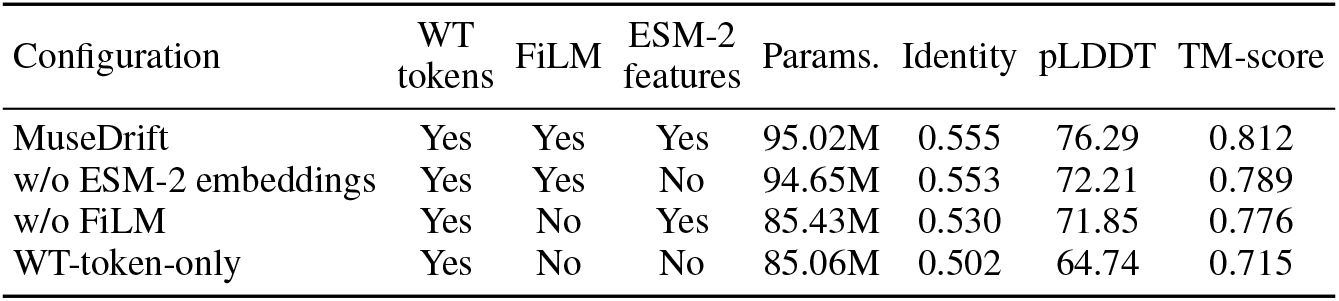
Comparison of conditioning configurations at nominal identity. .55. Equal-WT means over all 438 CAMEO WTs for the predesignated first sample. All configurations retain WT tokens; FiLM supplies numerical identity and ESM-2 supplies frozen WT features. Parameters exclude the frozen ESM-2 encoder. Training schedules differ, and configurations without FiLM use calibrated sampling temperatures; this is not a matched retraining ablation.

| Configuration | WT tokens | FiLM | ESM-2 features | Params. | Identity | pLDDT | TM-score |
| --- | --- | --- | --- | --- | --- | --- | --- |
| MuseDrift | Yes | Yes | Yes | 95.02M | 0.555 | 76.29 | 0.812 |
| w/o ESM-2 embeddings | Yes | Yes | No | 94.65M | 0.553 | 72.21 | 0.789 |
| w/o FiLM | Yes | No | Yes | 85.43M | 0.530 | 71.85 | 0.776 |
| WT-token-only | Yes | No | No | 85.06M | 0.502 | 64.74 | 0.715 |

**Table 22:** Realized identities for the Table 1 configuration comparison. Same first-sample, equal-WT aggregation as the structural scores.

| Configuration | .30 | .40 | .55 | .75 | .95 |
| --- | --- | --- | --- | --- | --- |
| MuseDrift | 0.316 | 0.407 | 0.555 | 0.755 | 0.949 |
| WT-token-only | 0.292 | 0.386 | 0.502 | 0.692 | 0.914 |

At the .55 setting, the full model has the highest pLDDT and predicted-WT-relative TM-score. The configuration without ESM-2 embeddings has pLDDT 72.21 versus 76.29 for the full model, at similar mean identity (.553 versus .555). The configuration without FiLM has pLDDT 71.85 and mean identity .530; the WT-token-only configuration gives pLDDT 64.74 and mean identity .502.

These results compare existing trained configurations rather than a controlled retraining study. The runs differ in effective batch size, learning-rate floor, gradient clipping, and selected checkpoint step. The temperature-controlled configurations also use calibration data and do not achieve the same identities as the directly conditioned models. The full configuration performs best in this comparison, but the differences do not isolate each component’s causal contribution.

### Temperature calibration for configurations without FiLM

The two configurations without FiLM use separate calibration curves on the same 60 frozen WTs: 30 CAMEO and 30 Mol-Instructions references from the released ProDVa panels (Liu et al., 2025). Records of length *L* ≤ 511 are shuffled with a shared Python `random.Random(2027)` stream, processing CAMEO first, and the first 30 per source are retained (manifest SHA-256 prefix `098d4452927d`). Each curve uses four samples per WT at 13 temperatures, 0.8, 0.9, …, 2.0, with identities averaged within WT, then equally across WTs. The curves decrease monotonically; piecewise-linear inversion gives the temperatures in Table 23, without endpoint clipping. Calibration does not update model weights. The full model and the FiLM configuration without ESM-2 features receive identity directly and do not use these mappings.

**Table 23:** Calibrated temperatures for configurations without FiLM. (four decimals). The configuration with ESM-2 features uses the displayed precision.

| Configuration | $\tau = .30$ | .40 | .55 | .75 | .95 |
| --- | --- | --- | --- | --- | --- |
| WT-token-only | 1.6913 | 1.4273 | 1.2557 | 1.0907 | 0.8844 |
| No FiLM (with ESM-2 features) | 1.8276 | 1.5721 | 1.3758 | 1.1963 | 0.9518 |

### Calibration references

The full 438-WT benchmark includes the 30 CAMEO references excluded in Appendix N’s 408-WT analysis. Their prepared-panel `cameo` suffixes are 00182, 00227, 00471, 00308, 00121, 00103, 00397, 00378, 00215, 00395, 00217, 00465, 00261, 00334, 00437, 00029, 00257, 00283, 00156, 00316, 00276, 00311, 00341, 00468, 00246, 00490, 00143, 00279, 00373, 00474. The `molinst` suffixes are 01426, 02692, 01412, 03619, 01337, 00113, 01343, 02822, 02409, 05333, 02844, 03090, 03032, 03767, 04760, 01894, 00262, 02393, 01347, 02085, 01870, 05069, 04472, 04558, 03112, 01141, 03244, 00398, 00868, 02424.

## R Scope of description-conditioned ProDVa generation

### Reference panel and comparison scope

We evaluate the released ProDVa generator on the fixed 429-WT subset in Appendix O, using function descriptions rather than WT sequences or target identities. Table 1 compares WT-conditioned generators on the full 438-WT panel. ProDVa’s outputs have sparse coverage near the requested identities used in that comparison, so we report its generation protocol and coverage here without a band-wise performance table.

### Description-conditioned ProDVa

We run the released ProDVa CAMEO checkpoint and official inference implementation (Liu et al., 2025). Inputs are the released function descriptions; the generator does not receive the WT sequence, WT length, or target identity. Retrieval uses the released training index and 16 descriptions per query. Before generation, the retrieval encoder is checked against 24 fixed stored vectors. Sampling uses temperature .7, top-*k* = 950, and a limit of 256 new tokens, with three predetermined seeds per reference record. There are 432 records for the 429 unique WTs, yielding 1,296 draws. All terminate with EOS; 107 are empty or contain nonstandard residues. We retain every draw in coverage accounting and do not resample or select by identity, length, or predicted quality.

### Realized identity and coverage

ProDVa can change sequence length. We therefore measure identity by global alignment using BLOSUM62, gap-open score −10, and gap-extension score −.5, penalizing terminal gaps. We use the first optimal alignment and divide identical residue pairs by all alignment columns, including gaps. The prespecified identity bands are [.25, .35), [.35, .45), [.50, .60), [.70, .80), and [.90, 1]. Only 73 of the 1,296 draws fall in these bands, of which 70 are valid standard-amino-acid sequences and receive pLDDT scores. The five bands contain 21, 4, 2, 1, and 2 WTs with scoreable outputs, respectively. These small, differing WT subsets do not support a paired comparison with the target-conditioned results in Table 1. All draws remain in the coverage accounting; no resampling or quality-based selection is applied.

### Structural scoring and overall outputs

All valid outputs are passed to the same single-sequence ESMFold checkpoint and scoring implementation as the WT-conditioned comparison: four configured recycles, chunk size 128, and a half-precision ESM tower. TM-align compares predicted variant and WT structures, normalized by WT length. WT structures are reused. No residues are deleted to make an output scoreable, and sequences shorter than three residues receive no TM-score. Across all identities, 1,189 outputs from 419 WTs have pLDDT scores; 1,100 outputs from 397 WTs have TM-scores. The 107 invalid outputs remain in the total draw count, and there are no folding failures. Scores are averaged over draws within a reference record, records within a WT, and then equally over WTs with the relevant metric. Overall mean pLDDT is 74.39 and mean TM-score is .322; mean WT pLDDT on the 429-reference panel is 85.34. These overall scores describe outputs spanning different lengths and distances, rather than a comparison at a matched WT identity, and do not establish retained function.

